# Bayesian inference of fitness landscapes via tree-structured branching processes

**DOI:** 10.1101/2025.01.24.634649

**Authors:** Xiang Ge Luo, Jack Kuipers, Kevin Rupp, Koichi Takahashi, Niko Beerenwinkel

## Abstract

**Motivation:** The complex dynamics of cancer evolution, driven by mutation and selection, underlies the molecular heterogeneity observed in tumors. The evolutionary histories of tumors of different patients can be encoded as mutation trees and reconstructed in high resolution from single-cell sequencing data, offering crucial insights for studying fitness effects of and epistasis among mutations. Existing models, however, either fail to separate mutation and selection or neglect the evolutionary histories encoded by the tumor phylogenetic trees.

**Results:** We introduce FiTree, a tree-structured multi-type branching process model with epistatic fitness parameterization and a Bayesian inference scheme to learn fitness landscapes from single-cell tumor mutation trees. Through simulations, we demonstrate that FiTree outperforms state-of-the-art methods in inferring the fitness landscape underlying tumor evolution. Applying FiTree to a single-cell acute myeloid leukemia dataset, we identify epistatic fitness effects consistent with known biological findings and quantify uncertainty in predicting future mutational events. The new model unifies probabilistic graphical models of cancer progression with population genetics, offering a principled framework for understanding tumor evolution and informing therapeutic strategies.

**Availability and implementation:** The `Python` package `FiTree` and the analysis workflows are available at https://github.com/cbg-ethz/FiTree.

## 1. Introduction

Cancer progression involves a somatic evolutionary process, where cells accumulate mutations that confer selective advantages, allowing them to evade immune surveillance and proliferate uncontrollably [Nowell, 1976, Hanahan and Weinberg, 2011, Greaves and Maley, 2012]. The extensive genetic and phenotypic heterogeneity within and between tumors enables cancer cells to rapidly adapt to different microenvironments, promoting metastasis and drug resistance [McGranahan and Swanton, 2017, Dagogo-Jack and Shaw, 2018]. To better understand the disease and develop effective treatment strategies, it is crucial to disentangle different evolutionary forces that shape the diverse tumor populations and improve evolutionary-aware predictions [Lässig et al., 2017, Watson et al., 2020, Salehi et al., 2021].

With the increasing quality and availability of single-cell DNA sequencing data for cancer research [Morita et al., 2020, Miles et al., 2020, Minussi et al., 2021, Zhang et al., 2023], we are now able to reconstruct the evolutionary histories of tumors at unprecedented resolution using specifically designed phylogenetic methods [e.g., Jahn et al., 2016, Zafar et al., 2019, Sollier et al., 2023, Sashittal et al., 2023, Kuipers et al., 2025]. Analyzing tumor phylogenies across patients has been successful in identifying reproducible features of cancer evolution, including common mutation orders and partitions [Ciccolella et al., 2021, Jahn et al., 2021], conserved evolutionary trajectories [Caravagna et al., 2018, Khakabimamaghani et al., 2019, Christensen et al., 2020, Hodzic et al., 2020, Pellegrina and Vandin, 2022], and patterns of co-occurrence and exclusivity among mutations [Kuipers et al., 2021, Luo et al., 2023, Ivanovic and El-Kebir, 2023]. These features facilitate predictions of tumor evolution with probabilistic measures and improve patient clustering for precise clinical diagnosis and prognosis [Baciu-Drăgan and Beerenwinkel, 2024]. However, most existing approaches for modeling intra-tumor phylogenies focus exclusively on the tree topologies, overlooking the abundances of different subclonal populations. Given that the frequencies of genetic variants are crucial markers for clinical disease classification and treatment planning [Bastian, 2014, Lee et al., 2019, Döhner et al., 2022], incorporating this information into the modeling framework is essential for uncovering the complex evolutionary dynamics of tumors and translating them into actionable insights for clinical applications.

When tree structures are not considered, population genetics models provide well-established tools to study how mutation, selection, and drift interact to shape the genetic composition and evolutionary trajectories of tumor cell populations over time [Ewens, 2004, Durrett, 2008, Beerenwinkel et al., 2015]. State-of-the-art methods, built upon the coalescent [Dinh et al., 2020, Werner et al., 2020], the Wright-Fisher process [Salehi et al., 2021], and branching processes [Williams et al., 2018, Dinh et al., 2020, Lang et al., 2020, Salichos et al., 2020, Lee and Bozic, 2022], have proven effective in inferring key parameters of cancer evolution (*e*.*g*. mutation rates and fitness advantages) from next-generation sequencing data. However, none of them can leverage the evolutionary trajectories encoded in intratumor phylogenetic trees to inform evolutionary parameter estimates. Some methods are tailored to bulk sequencing data [Williams et al., 2018, Dinh et al., 2020, Watson et al., 2020, Salichos et al., 2020], while others require time series measurements [Salehi et al., 2021, Lee and Bozic, 2022], which are not typically available from observational human data. Furthermore, these methods typically assume simple fitness landscapes [Wright, 1932], where fitness advantages are either uniform across mutations or independent of one another, neglecting the well-established effects of epistasis [Phillips, 2008, De Visser and Krug, 2014]. SCIFIL [Skums et al., 2019] is the first approach that considers both evolutionary histories and tumor cell abundances by superimposing a replicator model onto a tumor mutation tree—an alternative representation of tumor phylogenies [Jahn et al., 2016]. Yet, fitting a deterministic model to relative allele frequencies from a single patient inevitably leads to monotonically increasing fitness advantages toward the leaves of the tree. Therefore, SCIFIL is limited for inferring more complex fitness landscapes across patients. A more recent computational method infers the average fitness advantage of driver mutations by comparing simulated clone size vectors to the observed ones from multiple trees [Poon et al., 2025]. Despite its improvements, this approach disregards potential epistatic interactions and is computationally expensive due to simulation-based inference.

In this paper, we present FiTree, a tree-structured branching process model for learning fitness landscapes from single-cell tumor mutation trees. FiTree uniquely reconciles probabilistic graphical models of cancer progression with classical population genetics, providing a unified framework to analyze single-time-point intra-tumor data. In particular, we develop novel theory, extending the linear multitype branching processes [Athreya and Ney, 2004, Durrett and Moseley, 2010, Nicholson et al., 2023, Zhang and Bozic, 2024] to include a static initial wild-type population, tree structures, and a tumor-size-dependent censoring mechanism. We parametrize fitness landscapes with epistatic effects and leverage a Bayesian inference scheme to incorporate prior biological knowledge and quantify uncertainty in fitness estimates. Through simulations, we validate our branching process theory and demonstrate that FiTree outperforms state-of-the-art methods in recovering fitness landscapes. Finally, we apply FiTree to a single-cell acute myeloid leukemia (AML) dataset [Morita et al., 2020]. With the inferred posterior distribution of the fitness landscape, we provide uncertainty estimates of future events captured in longitudinal samples.

## 2. Materials and methods

FiTree is formulated as a probabilistic generative model for single-cell tumor mutation trees. The input to FiTree is a set of trees derived from independent patient samples (Fig. 1ab), and the output is the posterior distribution of the underlying fitness landscape (Fig. 1d). We begin by outlining the model setup, followed by the derivation of tree probabilities. Lastly, we describe the Bayesian inference scheme for parameter estimation. For reference, we summarize all key notations in Supp. Tab. S1.

**Fig. 1.**
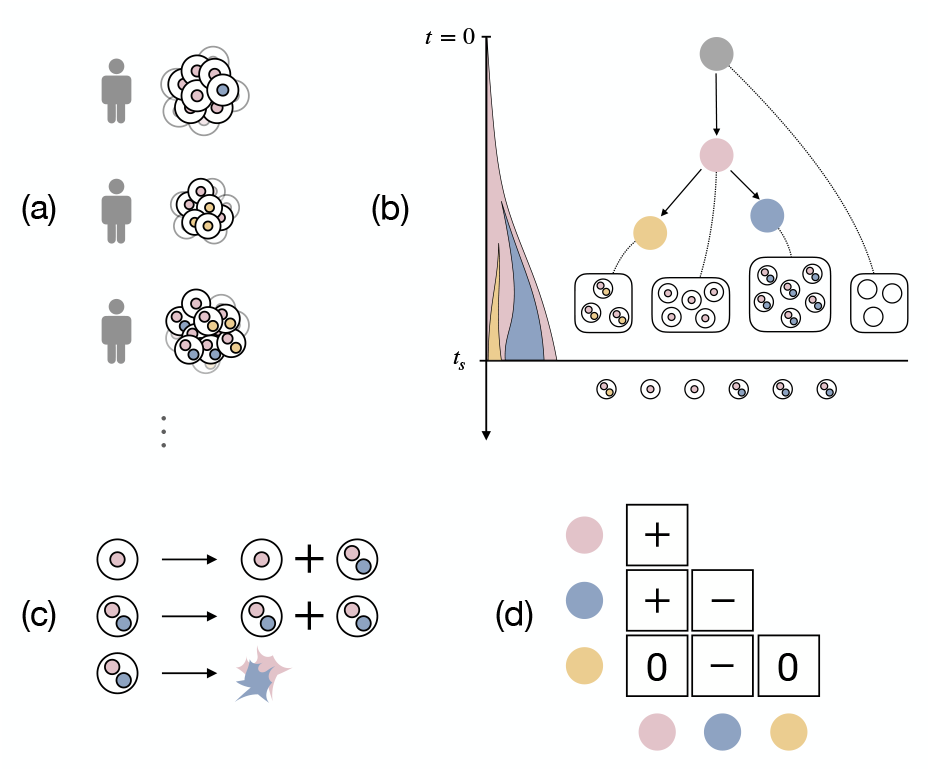
Overview of FiTree. (a) Cross-sectional single-cell data. Genetically diverse tumor cells of different patients are collected at single time-points. (b) Tumor evolutionary history represented as a tumor mutation tree and modeled using a probabilistic generative process in FiTree. The root node in gray corresponds to a static wild-type population at the start of evolution (*t* = 0). The colored nodes represent mutations accumulated over time. The sampling event occurs with rate proportional to the tumor size. At sampling time *t*_*s*_, single cells are collected for sequencing. (c) Illustration of the growth dynamics in the branching process, where cells can mutate, divide, or die stochastically. (d) A fitness matrix where diagonal entries represent fitness effects of individual mutations and off-diagonal entries correspond to the epistatic effects.

### 2.1. FiTree: model setup

A tumor mutation tree 𝒥 is a rooted tree that represents the evolutionary history of a tumor [Jahn et al., 2016] (Fig. 1b). Each node *v* in 𝒥 contains a subset of mutations *π*_*v*_ ⊆ [*n*] = {1, 2, …, *n*}. The cells attached to *v* form a subclone with genotype *g*_*v*_, determined by the mutations accumulated along the trajectory leading to *v*. The root node *v*_0_ corresponds to wild-type cells with no mutations. We denote the number of cells attached to *v* at time *t* by *C*_*v*_(*t*). To generate a tree 𝒥, we consider a tree-structured multi-type branching process that starts with a static population of wild-type cells, 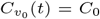 for all *t* ≥ 0, and no mutant cells. Namely, the tree initially consists of only the root, representing stable populations such as hematopoietic stem cells in bone marrow [Catlin et al., 2011] or intestinal stem cells in colorectal crypts [Kozar et al., 2013]. Then, a tree expansion rule *R* defines the set of subclones that could emerge next. One example of *R* is the Infinite Sites Assumption (ISA) [Kimura, 1969], which only allows mutations that have not yet occurred in any existing subclone to appear in the new subclones. Other rules relax ISA, permitting parallel, repeated, or back mutations [Kuipers et al., 2017]. Note that while the input trees may follow tree expansion rules imposed by upstream phylogenetic methods or expert knowledge, our model is not restricted to any specific rules, thereby providing a flexible framework for different applications.

We define the growth dynamics for cells within a mutant subclone *v* ≠ *v*_0_ as follows (Fig. 1c): assuming each cell grows independently of each other, cells of subclone *v* emerge from the parent subclone pa(*v*) through mutations at rate ν_*v*_, duplicate at rate *α*_*v*_, and die off at rate *β*_*v*_,

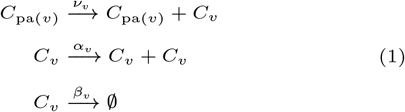

We introduce a sampling event *S*, which occurs at a rate proportional to the total number of mutant cells 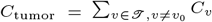,

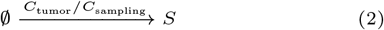

where *C*_sampling_ is a scaling factor representing the tumor size at which sampling is likely to occur. For instance, setting *C*_sampling_ = 10^11^ implies that, on average, a tumor is sampled when it reaches around 10^11^ cells. The process terminates either at the first arrival of *S* with sampling time *t*_*s*_, or upon reaching a pre-specified maximum observation time *t*_max_. The underlying principle is that larger tumors are more likely to be sampled, as patients are more prone to exhibit phenotypic symptoms and seek medical attention. Some patients, however, may develop cancer late in life and never get diagnosed, *i*.*e. t*_*s*_ *> t*_max_. In such cases, their tumor mutation trees are unobservable, introducing selection bias that must be accounted for during inference. Finally, to mimic the sequencing step, we sample *C*_seq_ cells without replacement from the tumor cell population. Typically the number of sequenced cells *C*_seq_ is much smaller than the total number of tumor cells *C*_tumor_ and often varies by patient. Leaf subclones with zero sequenced cells are considered unobserved and are therefore removed, resulting in the observed tumor mutation tree 𝒥.

Assuming that the rates of individual mutations, denoted by 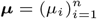, are independent and stay constant throughout the evolution, the subclone mutation rate is defined as 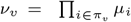, a product of the rates of the new mutations in *v*. For identifiability we assign a common death rate *β* to all cell types and allow the birth rates to vary. To capture both the fitness effects of individual mutations and potential epistasis, we parameterize the log birth-to-death ratio using a triangular fitness matrix *F* = (*f*_*ij*_)_*i*,*j*∈[*n*]_ ∈ ℝ ^*n×n*^ (Fig. 1d),

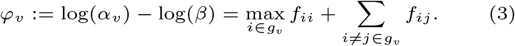

Here, the diagonal elements *f*_*ii*_ represent the fitness contributions of individual mutations, and the off-diagonal elements *f*_*ij*_ account for the pairwise interactions between mutations. This formulation assumes that only the most advantageous mutation dominates the fitness of a subclone, while all epistatic effects are captured in the interaction terms. The net growth rate is defined as

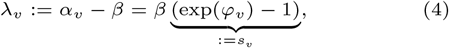

with its sign indicating whether the mutations are collectively advantageous (*λ*_*v*_ *>* 0), neutral (*λ*_*v*_ = 0), or deleterious (*λ*_*v*_ *<* 0). By definition the wild-type cells have a net growth rate of zero, *i*.*e*. 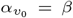 and 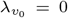. Hence, the term *s*_*v*_ can be interpreted as the relative increase from the wild-type division rate.

### 2.2 Calculation of tree probabilities

To compute the probability of observing a tumor mutation tree 𝒥 at sampling time *t*_*s*_, we first need to understand the subclonal population and sampling time distributions. However, finding the exact solution of the above system is a highly intricate problem [Athreya and Ney, 2004, Antal and Krapivsky, 2011], even under the assumption that all data and parameters are perfectly known, which will be relaxed in the later sections. Therefore, we resort to approximations in the limit of large times and small mutation rates [Durrett and Moseley, 2010, Nicholson et al., 2023], which is often a reasonable assumption in cancer studies. In particular, we generalize the results in Nicholson et al. [2023], which considers a linear multi-type branching process initiating from a single cell that grows exponentially, to model tree structures with static initial wild-type populations followed by cell types of arbitrary fitness. We show in Theorem 1 (Supp. Sec. B.1) that for any mutant type *v ≠ v*_0_ in 𝒥, the limiting distribution of the subclonal population *C*_*v*_(*t*) can be decomposed approximately into a time-independent random variable 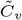 and a time-dependent deterministic function,

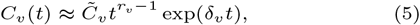

where *r*_*v*_ is the number of times the running-max net growth rate

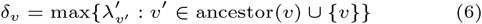

is attained along the lineage of *v*. Intuitively, given the subclonal population sizes

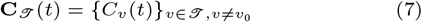

at some large time *t*, this result enables us to approximate deterministically the unobserved evolutionary trajectories **C**_*T*_ (*s*) for all 0 *< s < t*, thereby significantly simplifying the calculation of probabilities.

At any given time *t*, the specific size distribution of a subclone *v* is determined by both its position in the tree and its net growth rate *λ*_*v*_. If *v* comes directly after the root, *i*.*e*. pa(*v*) = *v*_0_, then its size *C*_*v*_(*t*) follows a negative binomial distribution, which in the large time limit is decomposable as in Eq. (5),

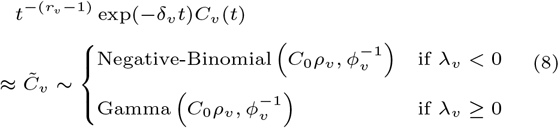

where

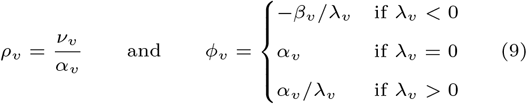

are the shape and scale, respectively, of the distribution (Lemma 1 & Theorem 1, Supp. Sec. B.1).

For all remaining subclones *v* with pa(*v*) ≠ *v*_0_, the population distribution has no simple form. It is characterized by the Laplace transform conditioned on the parent subclone size,

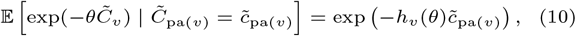

where *h*_*v*_(θ) is a piece-wise function that distinguishes whether *v* is fitter (*λ*_*v*_ *>* δ_pa(*v*)_), equally fit (*λ*_*v*_ = δ_pa(*v*)_), or less fit (*λ*_*v*_ *<* δ_pa(*v*)_) than all its ancestor subclones (Theorem 2, Supp. Sec. B.1). If *v* is not fitter than its parent, then on average, its size will remain proportional to that of its parent at large times. Otherwise, it expands at its own net growth rate. In other words, the growth dynamics of subclone *v* are dominated by the fittest subclone in its lineage. Thus, deleterious mutations can still persist in the cohort by hitchhiking alongside beneficial mutations already present in the subclone.

The sampling time *T*_*s*_ is the waiting time until the first arrival of the sampling event *S*(*t*). By definition it follows a nonhomogeneous Poisson process with intensity function depending on the tumor size,

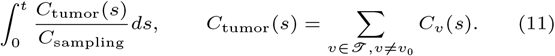

Hence, as proved in Theorem 3 (Supp. Sec. B.2), the distribution function of *T*_*s*_ is given by

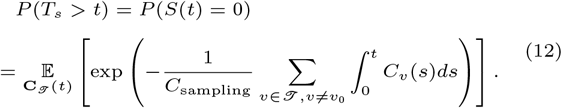

In addition, the vector of sequenced cells, defined as

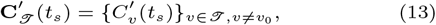

are sampled without replacement from the subclonal population size vector (Eq. (7)), thereby following a multivariate hypergeometric distribution with density

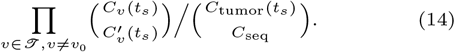

Putting everything together, the probability of observing a tumor mutation tree 𝒥 at sampling time *t*_*s*_ is given by

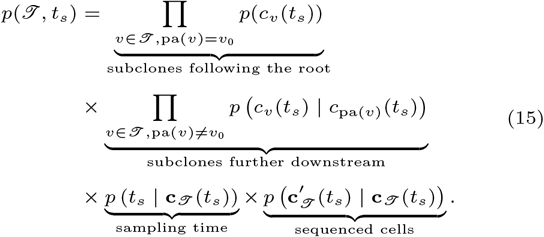

We refer to this probabilistic model of mutation trees as FiTree.

### 2.3. Bayesian inference of fitness landscapes

The parameters of FiTree (Eq. 15) are

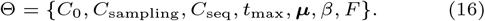

In this paper, we focus on inferring the marginal posterior distribution of the fitness matrix *F* given a set of *N* tumor mutation trees 𝒥 = *{* 𝒥 ^*j*^ *}*_*j*∈*N*_ with sampling times 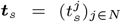. To this end, we adopt the following assumptions about the data and the parameters:

1. While the actual subclonal population sizes (Eq. (7)) are unobservable, the total tumor size *C*_tumor_ can be estimated from clinical measurements.
2. The sample proportions accurately reflect the true cancer cell fractions in patients,

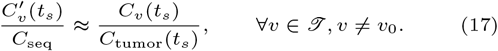
3. All parameters, except *F*, can be derived from either clinical data or literature (see Supp. Sec. E.1 for an example).

Note that the sampling times ***t***_*s*_, if unavailable, can still be readily integrated out, which is not a limiting factor for broad applications. The posterior of *F* is given by

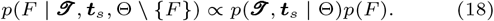

Here, the log-likelihood *p*(𝒥, ***t***_*s*_ | Θ) is computed as

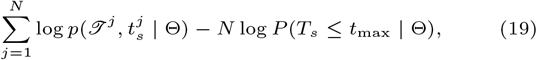

where the first term sums the log-likelihoods of all trees (Eq. (15)), and the second term adjusts for selection bias, as datasets can only contain observable trees and no negative controls. When the lifetime risk of developing and being diagnosed with the disease,

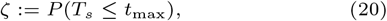

can be reliably estimated from external databases [e.g., Surveillance, Epidemiology, and End Results (SEER) Program], we can additionally impose a prior on *P* (*T*_*s*_ ≤ *t*_max_ | Θ), centered at ζ with a small width, to enforce consistency. Moreover, we assign the normal prior 𝒩 (0, *σ*^2^) to every distinct entry in the matrix. To explore the parameter space, we use Markov chain Monte Carlo sampling. Further details are given in Supp. Sec. C.

## 3. Results

### 3.1. Simulations

Our simulation studies consist of two parts: (1) validating the large-time approximations for subclonal population and sampling time distributions, and (2) benchmarking the performance of FiTree in learning the subclone fitness effects φ_*v*_ (Eq. (3)) from simulated tumor mutation trees against state-of-the-art methods.

#### 3.1.1. Validating approximated distribution functions

We begin with verifying the large-time approximations of the subclonal population and sampling time distributions, which are the building blocks of the likelihood function (Eq. (15)). Specifically, we perform stochastic simulations of branching processes with three subclones and a sampling event using the Gillespie algorithm with tau-leaping [Gillespie, 2001]. The simulations encompass seven different parameter settings, as summarized in Supp. Tab. S2 and illustrated in Fig. 2a. While most parameters follow those in [Nicholson et al., 2023], covering a range of net growth rates and mutation rates, we extend the analysis to include neutral and deleterious cases for the first cell type, as well as linear versus branching structures. Further simulation details are provided in Supp. Sec. D.1. Across all scenarios (Fig. 2b), the empirical marginal cumulative distribution functions (CDF) for each cell type and sampling time closely match their analytical approximations (Corollary 1 and Theorem 3, Supp. Sec. B). Accurate approximations of the distribution functions ensure reliable fitness landscape inference in downstream analysis.

**Fig. 2.**
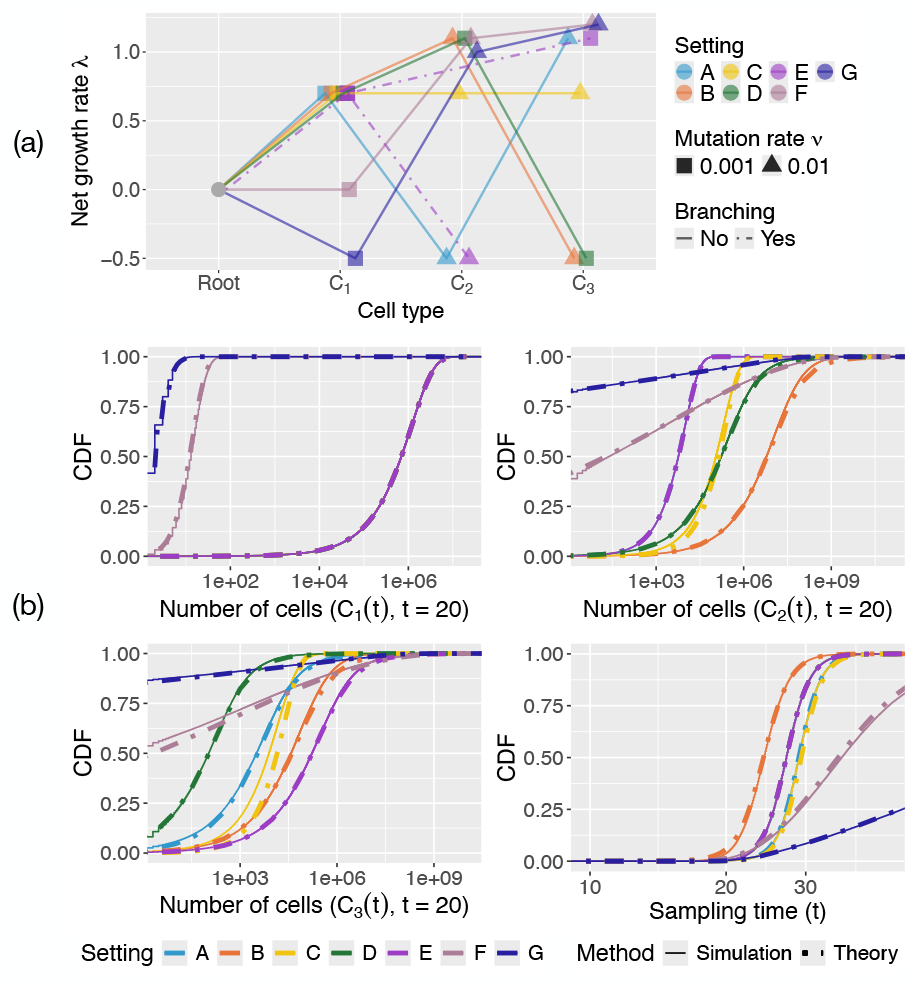
Accurate approximations of subclonal population and sampling time distributions in tree-structured branching processes. (a) Seven parameter settings used in stochastic simulations. The *x*-axis shows cell types; the *y*-axis shows net growth rates. Linear structures (solid lines) are used except for setting E (branching structure, dashed lines). Shapes indicate mutation rates (rectangle: 0.001; triangle: 0.01). The root has net growth rate 0 and no mutation rate. (b) Comparison of cumulative distribution functions from simulations (solid) and theoretical approximations (dashed). Some curves overlap, *e*.*g*. settings A–E for *C*_1_(*t*), as plots are faceted by variables and colored by settings. Alternate views faceted by settings are in Supp. Fig. S2–S8.

#### 3.1.2. Benchmarking FiTree performance

The second part of the simulations benchmarks FiTree’s performance in learning fitness landscapes from tumor mutation trees against state-of-the-art methods, including SCIFIL [Skums et al., 2019], fitclone [Salehi et al., 2021], and the diffusion approximation of a one-type branching process [Watson et al., 2020]. The trees are simulated using the Gillespie algorithm with tau-leaping from the tree-structured multitype branching process defined in Section 2.1. The pseudocode for tree generation, along with details on ground-truth parameter selection and method execution, are provided in Supp. Sec. D.

Since fitness definitions differ across models, direct comparisons of absolute fitness effects are generally not possible, except for the method in Watson et al. [2020]. For this method, we separately compare the deviation between the subclone fitness estimates 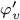 and the ground truth φ_*v*_ using the following metrics:

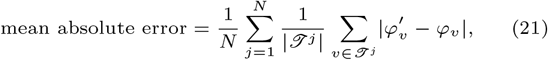

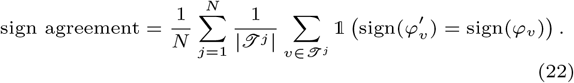

For all methods, we evaluate Spearman correlations between ground-truth and estimated fitnesses to measure each model’s ability to capture relative variations in fitness landscapes. Here, we include a simple baseline that ranks subclone fitnesses based on their average frequencies within the cohort. Since SCIFIL and fitclone are designed for single-patient analyses, we apply them first on individual trees and average the fitness effects across all trees. To evaluate generalizability, we assess how well each method estimates the fitness of both observed and unobserved subclones. Observed subclones are those with non-zero cell counts in the trees. Unobserved subclones are biologically plausible subclones with zero sequenced cells. They could have emerged but are not present in the trees, potentially due to late emergence or technological detection limits. Specifically, we augment the true trees by attaching empty subclones and focus on genotypes with up to two mutations, as including more provides little additional insight given that none of the methods capture higher-order epistasis. This augmentation does not affect the structure or interpretation of the trees, nor does it bias fitness estimates, but allows all methods to provide fitness estimates for both observed and unobserved subclones.

For varying number of mutations (*n* ∈ *{*5, 10, 15*}*) and number of samples (*N* ∈ *{*200, 500*}*), FiTree consistently outperforms all alternative methods in learning the fitness values of both observed and unobserved subclones (Fig. 3). With additional experiments, we show that removing less frequently mutated genes from inference does not create strong bias on the estimated fitness landscape (Supp. Fig. S12) and that imposing a strong informative prior on the lifetime risk (Eq. (20)) mitigates overestimation of fitness effects (Supp. Fig. S13).

**Fig. 3.**
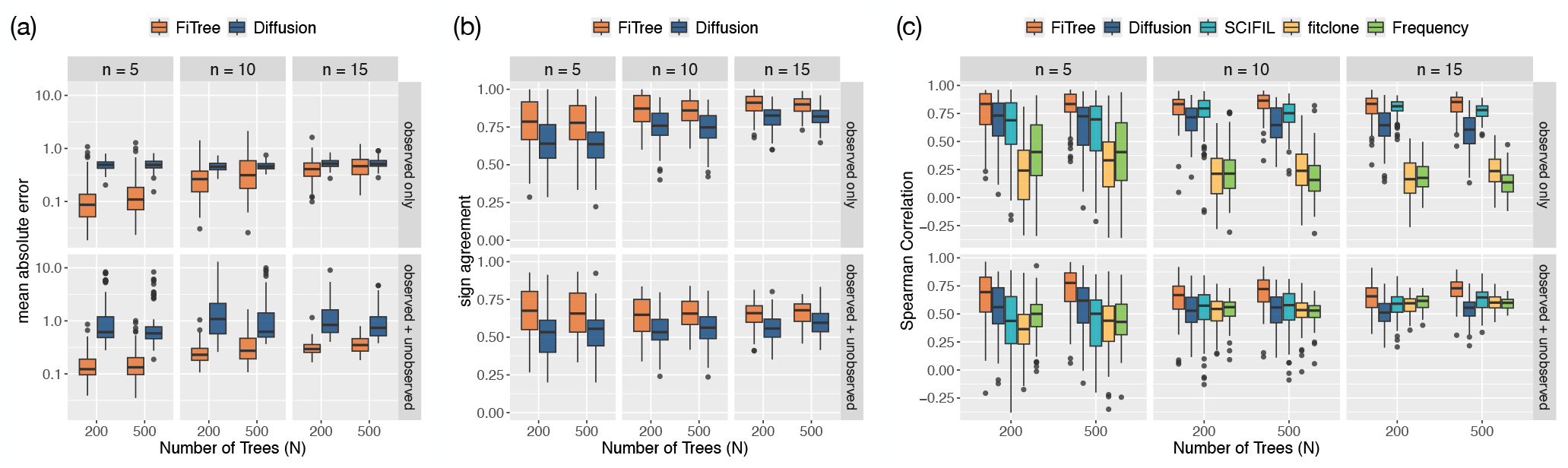
Performance comparison of FiTree with state-of-the-art methods in learning fitness landscapes on simulated data. Boxplots display the mean absolute errors (a), sign agreement (b), and Spearman correlations (c) between inferred and ground truth fitness values across 100 simulation runs, grouped by *n* ∈ {5, 10, 15}, *N* ∈ {200, 500}, and evaluation scope (observed subclones only or including unobserved subclones). For FiTree and fitclone, each point is derived from the posterior medians, while for other methods, points correspond to maximum likelihood estimates. Boxes represent the interquartile range (IQR) with medians marked inside, whiskers extend to 1.5 times the IQR, and outliers are shown as individual points.

The diffusion approximation approach performs reasonably well, as it is also based on a branching process model, albeit a much simpler one, and aggregates information across patients. However, being designed for bulk sequencing data, it cannot fully leverage the evolutionary trajectories and subclone sizes provided by single-cell tumor mutation trees. While its simplicity offers some stability as the problem dimensions increase (Fig. 3a), this comes at the cost of ignoring interactions between mutations, leaving little room for improvement with additional data (Fig. 3bc). In contrast, FiTree captures these interactions and provides a more comprehensive model of the evolutionary process, resulting in better performance despite requiring more data and computational effort, particularly in higher-dimensional settings.

As single-patient models, SCIFIL and fitclone assign fitness values to subclones based primarily on their sizes within individual trees. A subclone may have high fitness if it is large in one tree but low fitness if it is small or absent in another. While differences in tumor microenvironments across patients might lead to varying fitness for the same subclone, this does not fully explain the common patterns observed across trees in real data. A subclone may appear small in one tree not due to lower fitness but because of later emergence, reflecting the stochastic nature of the evolutionary process, which deterministic models like SCIFIL cannot capture. While SCIFIL performs well for observed subclones, likely due to its consideration of tree structures, its performance is similar to the baseline method when unobserved subclones are included (Fig. 3c).

Even as a stochastic method, fitclone struggles when it is provided with only two time points: a trivial initial point at *t* = 0, with no mutations present, and a second point at diagnosis, reflecting the limitations of many real-world datasets. The poor results suggest that fitclone requires more time points to be effective and cannot reliably handle such sparse data.

The runtime of FiTree scales with the number of samples or trees (*N*), the total number of genotypes across all trees (more precisely, the union tree size, see Supp. Sec. C.3), the number of fitness parameters 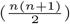, as well as the number of MCMC samples and chains used during inference. A comparison of runtimes is illustrated in Supp. Fig. S14.

### 3.2. Application to a single-cell AML dataset

We apply FiTree to a single-cell AML dataset [Morita et al., 2020] covering 531 mutations in *n* = 31 genes and comprising *N* = 123 tumor mutation trees, each from a different patient, reconstructed using SCITE [Jahn et al., 2016]. To address the challenge of high dimensionality with a limited number of samples and enhance the statistical power to detect non-random patterns, we summarize the variants at the gene level and focus on mutations in genotypes present in at least three patients. This filtering reduces the analysis to *n* = 15 genes, which were also identified by TreeMHN [Luo et al., 2023] as having statistically stable exclusivity patterns. Importantly, less frequently mutated genes are not removed from the trees but are excluded from the inference of epistatic fitness effects, with their corresponding off-diagonal entries in the fitness matrix *F* assigned zeros. As validated in simulations, this approach does not introduce significant bias in the estimates. We provide details on data preparation, including tumor size estimation, and static parameter estimation, such as mutation rates and other model parameters from Eq. (16), Supp. Sec. E.

The mutation rates and the posterior distribution of the fitness matrix inferred by FiTree are shown in Fig. 4a. The results reveal a complex interplay of positive and negative epistatic effects, suggesting that assuming uniform [Poon et al., 2025] or independent [Watson et al., 2020] fitness effects for individual mutations is insufficient to capture the underlying evolutionary dynamics. For comparison, fitness estimates from alternative methods and the corresponding runtimes are summarized in Supp. Tab. S4 and S5, respectively.

**Fig. 4.**
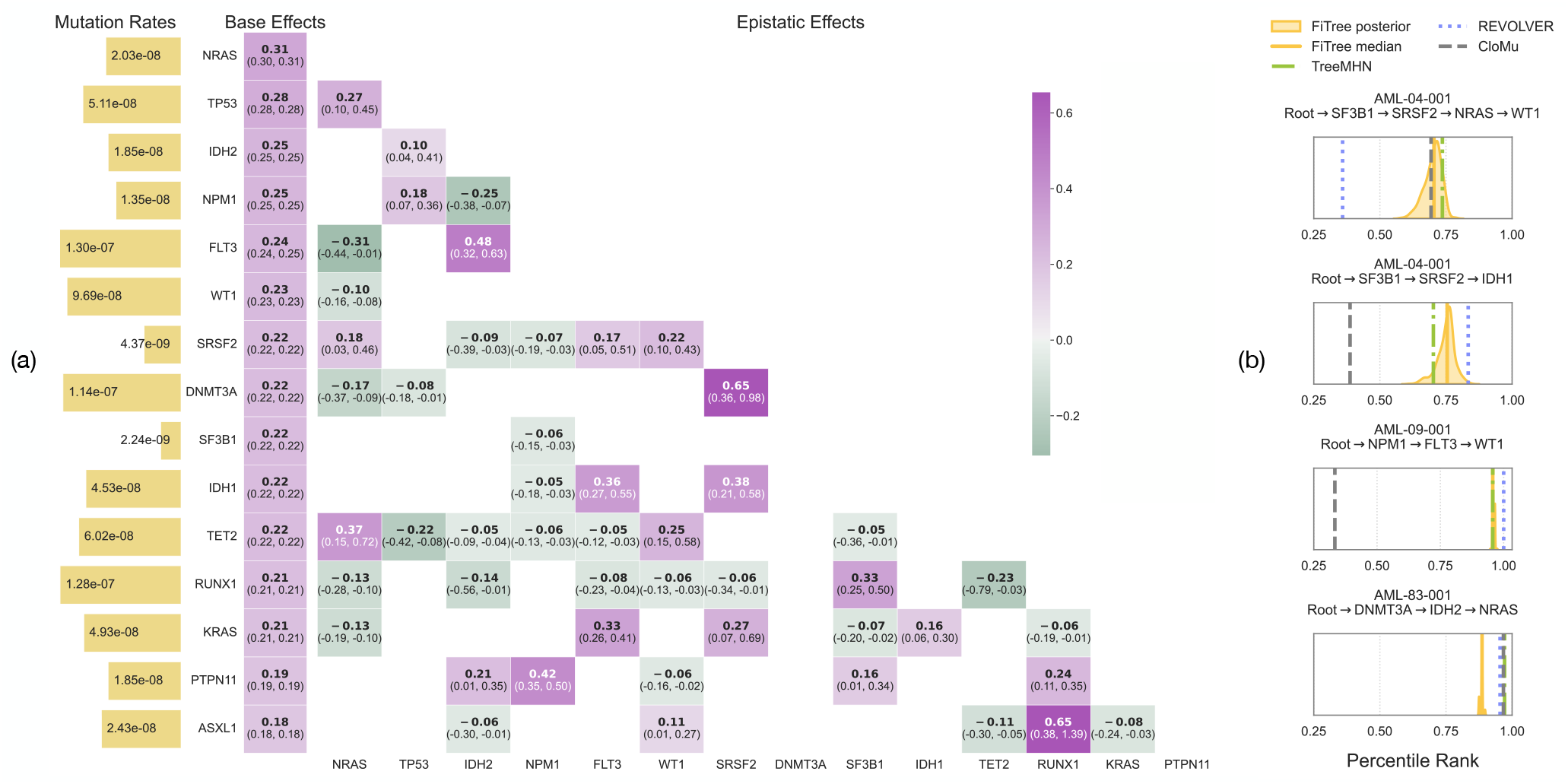
(a) Mutation rates and fitness matrix for the AML dataset. The mutation rates are displayed as bar plots on the left (yellow), with bar lengths scaled logarithmically according to the labeled mutation rates. The inferred fitness matrix is shown as a heatmap, where base effects are separately arranged in a column on the left. Matrix rows and columns are ordered by decreasing base effects. Each cell displays the posterior median in bold, with brackets indicating the corresponding 95% highest density interval (HDI). Cells with absolute epistatic effects lower than 0.05 or HDIs covering zero are masked. Full fitness matrix along with standard deviations are provided in Supp. Fig. S16 and S17. (b) Estimated percentile ranks of the four new events in the first post-treatment samples for patients AML-04-001, AML-09-001, and AML-83-001. Each subplot corresponds to one event. The density curves are estimated based on 1000 samples drawn from the FiTree posterior distribution. The lines represent the posterior median of FiTree (yellow, solid), as well as the maximum likelihood estimates by TreeMHN (green, dash-dotted), REVOLVER (blue, dotted), and CloMu (dashed).

All base fitness effects of the frequently mutated genes are positive, with increases relative to the wild-type division rate ranging from 20% to 36% (*s*_*v*_ in Eq. (4)), confirming their role as driver mutations that confer a significant growth advantage to tumor cells. *DNMT3A* and *FLT3*, two commonly mutated genes in AML [Miles et al., 2020, Morita et al., 2020], have similar mutation rates, yet *FLT3* exhibits higher base fitness. One plausible explanation is that mutations in epigenetic modifiers (*e*.*g. DNMT3A*) are likely initiating events [Dai et al., 2017], whereas those in signaling genes (*e*.*g. FLT3*) drive rapid tumor growth, resulting in diagnosis with fewer subsequent mutations [Miles et al., 2020]. In contrast, TreeMHN, which does not account for subclone sizes, estimates the highest fixation rate for *DNMT3A*. This does not contradict our findings; rather, it suggests that *DNMT3A* mutations are expected to appear more frequently in the cohort but do not necessarily result in larger subclones. By modeling mutation and selection separately while incorporating size information, FiTree provides more biologically meaningful insights.

With our Bayesian inference framework, we observe that the 95% highest density intervals for base effects are narrower than those for epistatic effects, indicating greater uncertainty in estimating epistasis due to the limited sample size. We observe large positive interactions with less uncertainty for several pairs of genes such as *NPM1* -*PTPN11, RUNX1* -*ASXL1*, and *FLT3* - *IDH1/2*. They correspond to known co-occurrence, and some are associated with worsened clinical outcomes [DiNardo et al., 2015, Falini et al., 2020, Gaidzik et al., 2016]. Additionally, FiTree identifies a negative epistatic effect between *FLT3* and *NRAS*, both of which exhibit high base fitness. While each mutation provides a strong selective advantage individually, their combined presence may result in reduced fitness, making them less likely to co-occur within the same genome. This finding aligns with the mutual exclusivity observed in previous studies [Kuipers et al., 2021, Luo et al., 2023] and offers an explicit evolutionary explanation for the reported secondary resistance to *FLT3* inhibitors, which may arise from off-target RAS pathway mutations (*e*.*g. NRAS*) present in small populations before treatment [McMahon et al., 2019].

By sampling from the posterior distribution of the fitness matrix, we assess FiTree’s ability to predict the next likely mutational events given a primary tumor tree. We validate our evolutionary forecasts using longitudinal samples from the same cohort, which include clinical data for three patients with four new mutational events (*i*.*e*. subclones with new genotypes) in the first post-treatment samples [Morita et al., 2020]. The predicted likelihood of an event is computed as the instantaneous growth rate, defined as the product of the parent subclone size at sampling, the mutation rate, and the estimated running-max growth rate (Eq. (6)). Following Luo et al. [2023], we rank the instantaneous growth rates among all possible future events for a given tree, where a higher percentile rank indicates a greater chance of occurrence. FiTree assigns high rankings to all four events (Fig. 4b), consistent with previous consensus estimates by TreeMHN, REVOLVER, and CloMu. Unlike point estimation methods, FiTree offers a probabilistic view that reveals the inherent complexity of the evolutionary process, and supplies credible intervals.

## 4. Discussion

We have introduced FiTree, a novel tree-structured branching process model with a Bayesian inference scheme, to infer complex fitness landscapes from single-cell tumor mutation trees. Through simulations, FiTree demonstrated improved performance in recovering fitness landscapes compared to state-of-the-art methods.

When applied to a single-cell AML dataset, FiTree identified epistatic fitness effects associated with known biological findings and demonstrated comparable performance to existing probabilistic models (TreeMHN, REVOLVER, CloMu) in predicting future mutational events in longitudinal samples. In addition, FiTree uniquely quantifies the uncertainty associated with fitness estimates and future mutations, offering valuable insights into tumor progression and enabling evolutionary forecasting. More accurate predictions will require additional data and explicit modeling of treatment effects, which can now be achieved with FiTree’s design. Specifically, treatment-induced changes can be incorporated into the fitness parameterization, enabling relapse simulation through forward sampling. Such analyses are not feasible with previous methods, which lack a growth model and an explicit notion of fitness.

Moreover, FiTree could be further developed to incorporate patient-specific factors, such as age, sex, pre-existing conditions (*e*.*g*. smoking history), tissue type, and cancer subtype, which can impact the underlying fitness landscapes of tumor evolution. Integrating these factors would result in a more comprehensive model, where personalized fitness estimates could support downstream analyses such as patient stratification, survival prediction, and treatment planning [Bayer et al., 2023].

FiTree is particularly well-suited for analyzing liquid cancers like AML, as branching processes are designed for homogeneously mixed populations with non-spatial growth [Durrett, 2015], and the assumption that sampled proportions accurately reflect the true tumor cell composition often holds. However, real-world data, especially from solid tumors, typically violate this assumption and introduce further complications from spatial heterogeneity, biased samples, limited cell numbers and sequencing errors, leading to large uncertainty in the mutation trees. Consequently, fitness values may be overestimated for genotypes that are sampled and underestimated for those that are not. FiTree can partially mitigate these biases by accepting multiple weighted trees per patient, such as those from different lesions or posterior sampling. Fully addressing tree uncertainty and spatial constraints remains an important direction for future work.

While this paper primarily focused on trees derived from DNA data, the model itself does not assume genetic events exclusively. In principle, the framework could also accommodate heritable non-genetic events, such as epigenetic modifications. Prior work applying branching process models to phenotypic switching [Gunnarsson et al., 2023] suggests that extending FiTree in this direction is possible, though it would require careful adaptation of both data types and model assumptions. FiTree’s parameter space increases quadratically with the number of mutations, demanding larger sample sizes for robust performance. For smaller cohorts, users can reduce model complexity by focusing on fewer mutations or grouping mutations by pathways to obtain higher-level fitness estimates.

The current parameterization considers only pairwise epistatic effects and constant mutation rates over time. These simplifications are practical when data are limited. However, real tumors can exhibit more complex interactions, such as clonal cooperation [Tabassum and Polyak, 2015]. Also, mutation rates may shift, for example, in response to genomic instability or treatment interventions. As more data becomes available, extending FiTree to capture higher-order epistasis and time-dependent mutation rates may yield a more accurate representation of tumor evolution and enhance predictive power.

In summary, FiTree provides a unified approach that bridges probabilistic graphical models of cancer progression with classical population genetics for analyzing single-cell data. It opens up opportunities for more precise modeling of tumor progression and drug response, enabling improved treatment strategies and personalized interventions.

## 6. Author contributions statement

Conceptualization: XL, JK, KR, NB; Data Curation: XL, KT; Formal Analysis: XL; Methodology: XL, JK, KR, NB; Software: XL; Supervision: JK, NB; Visualization: XL; Writing – Original Draft Preparation: XL; Writing – Review & Editing: all authors.

## 7. Acknowledgments

This work was supported by ETH Zurich [Open ETH project SKINTEGRITY.CH (N.B.)] and the European Union’s Horizon 2020 research and innovation program [grant number 951970, OLISSIPO project (N.B.)].

