## Supplementary Information for "Bayesian inference of fitness landscapes via tree-structured branching processes"

**References** **32**

### A Notations

| **Notation** | **Description** |
|---|---|
| $n$ | Number of mutations |
| $N$ | Number of samples |
| $\mathscr{T}$ | Tumor mutation tree |
| $T_s$ | Sampling time of the tumor, or the first arrival time of the sampling event $S$ |
| $v_0$ | Root node of tumor mutation trees, representing the wild-type population |
| $\pi_v \subseteq [n]$ | Set of mutations in node $v$ |
| $g_v \subseteq [n]$ | Genotype of $v$, which is the set of mutations accumulated along the trajectory from the root to the node where the subclone locates |
| $R$ | Tree expansion rule (*e.g.* infinite sites assumption) |
| $C_v(t)$ | The number of cells in $v$ at time $t$ |
| $C_{v_0}(t) = C_0$ | Wild-type compartment with a static population size $C_0$ as a scaling factor for the initial mutation rates (*e.g.* $10^8$ cells) |
| $C_{\text{tumor}}(t) = \sum_{v \in \mathscr{T}, v \neq v_0} C_v(t)$ | Total number of mutant cells in a tree $\mathscr{T}$ at time $t$ |
| $C_{\text{sampling}}$ | Scale of the number of cells at sampling (*e.g.* $10^9$ cells) |
| $C_{\text{seq}}$ | Number of cells being sequenced (*e.g.* $10^5$ cells) |
| $\text{pa}(v)$ | The parent subclone of $v$, which is unique |
| $\text{ch}(v)$ | The children set of subclones of $v$ |
| $\text{anc}(v)$ | The ancestor set of subclones of $v$ |
| $\text{mrca}(v)$ | The most recent common ancestor subclone of $v$, *i.e.* $v' \in \text{anc}(v) \cup \{v\}$ and $pa(v') = v_0 \iff v' = \text{mrca}(v)$ |
| $\boldsymbol{\mu} = (\mu_i)_{i \in [n]}$ | Mutation rates of individual mutational events |
| $\nu_v = \prod_{i \in \pi_v} \mu_i$ | Mutation rate of a subclone $v$, defined by the product of mutation rates of the mutations in $v$ |
| $F = (f_{ij})_{i,j \in [n]}$ | Fitness parameter matrix |
| $\alpha_v$ | Birth rate of cells in subclone $v$ |
| $\beta_v = \beta$ | Death rate of cells in subclone $v$, which is assumed to be the same for all subclones |
| $\lambda_v = \alpha_v - \beta_v$ | Net growth rate of cells in subclone $v$ |
| $\varphi_v = \log(\alpha_v) - \log(\beta_v)$ | Log birth-to-death ratio of subclone $v$ |
| $s_v = \exp(\varphi_v) - 1$ | Relative increase from the wild-type division rate $\beta$ |
| $\delta_v$ | Running-max net growth rate along the trajectory of $v$ |
| $r_v$ | Number of times the running-max net growth rate has been attained along the trajectory of $v$ |
| $\rho_v$ | Shape of the population distribution of subclone $v$ |
| $\phi_v$ | Scale of the population distribution of subclone $v$ |
| $\gamma_v$ | Growth ratio of subclone $v$ |

Table S1: A summary of key notations used in this article.

### B Population dynamics in a tree-structured branching process

#### B.1 Population distributions

In the following lemma, we will show that the number of cells in the mutant populations directly following the root can be decomposed into a product of a time-dependent deterministic function and a random variable independent of time.

**Lemma 1.** *Let $(X(t))_{t\geq 0}$ be a one-type branching process, where cells mutate from a static population of wild-type cells at rate $\nu > 0$, duplicate at rate $\alpha > 0$, and die at rate $\beta > 0$:*

$$\begin{aligned} \emptyset &\xrightarrow{\nu} X \\ X &\xrightarrow{\alpha} X + X \\ X &\xrightarrow{\beta} \emptyset \end{aligned} \tag{1}$$

*Suppose initially there are no cells, i.e. $X(0) = 0$. Let $\lambda = \alpha - \beta$ and $\rho = \frac{\nu}{\alpha}$. Then $X(t)$ has the following large-time limiting distributions:*

*(i) if $\lambda < 0$, then*

$$\lim_{t\to\infty} X(t) = Y \sim \textit{Negative-Binomial}\left(\rho, -\frac{\lambda}{\beta}\right); \tag{2}$$

*(ii) if $\lambda = 0$, then*

$$\lim_{t\to\infty} t^{-1}X(t) = Y \sim \textit{Gamma}\left(\rho, \frac{1}{\alpha}\right); \tag{3}$$

*(iii) if $\lambda > 0$, then*

$$\lim_{t\to\infty} \exp(-\lambda t)X(t) = Y \sim \textit{Gamma}\left(\rho, \frac{\lambda}{\alpha}\right). \tag{4}$$

*Proof.* We will begin with recapitulating a classical result for one-type branching processes, stating that $X(t)$ follows a negative binomial distribution. First, the probability generating function of $X(t)$ is

$$G(s,t) = \sum_{k=0}^{\infty} s^k P(X(t) = k). \tag{5}$$

The corresponding master equation is

$$\begin{aligned} \frac{dP(X(t) = k)}{dt} &= [\nu + \alpha(k-1)]P(X(t) = k-1) \\ &\quad + \beta(k+1)P(X(t) = k+1) \\ &\quad - [\nu + \alpha k + \beta k]P(X(t) = k). \end{aligned} \tag{6}$$

Multiplying both sides by $s^k$ and summing over $k = 0, 1, 2, \ldots$ give

$$G_t = [\alpha s(s-1) + \beta(1-s)]G_s + \nu(s-1)G, \tag{7}$$

where $G_t = \frac{\partial G(s,t)}{\partial t}$ and $G_s = \frac{\partial G(s,t)}{\partial s}$. Then, we apply the method of characteristics and solve the following system of differential equations

$$\begin{cases} \dfrac{ds}{dt} = \alpha s(s-1) + \beta(1-s) := \omega \\ \dfrac{dG}{dt} = \nu(s-1)G \\ G(s,0) = 1 \end{cases} \tag{8}$$

where the last line corresponds to the initial condition that we start with no cells. One can verify that

$$G(s(0),t)=\exp\left[\nu\int_0^t (s(t')-1)dt'\right] \tag{9}$$

where

$$s(t)-1=\begin{cases}\dfrac{s(0)-1}{1-\alpha t(s(0)-1)} & \text{if } \lambda=0,\\ \dfrac{\lambda(1-s(0))}{\alpha(s(0)-1)-(\alpha s(0)-\beta)\exp(-\lambda t)} & \text{otherwise.}\end{cases} \tag{10}$$

Note that we are solving $\frac{ds}{dt}=\omega$ instead of $\frac{ds}{dt}=-\omega$ as in the usual step of the method of characteristics, which allows us to directly replace $s(0)$ by $s(t)$ in the final expression of $G(s(0),t)$ after integration and obtain

$$G(s,t)=\begin{cases}\left[\dfrac{1}{1-\alpha t(s-1)}\right]^\rho & \text{if } \lambda=0,\\ \left[\dfrac{\lambda}{\alpha(s-1)-(\alpha s-\beta)\exp(\lambda t)}\right]^\rho & \text{otherwise,}\end{cases} \tag{11}$$

where $\rho=\frac{\nu}{\alpha}$. With the transformations

$$q(t)=\begin{cases}1+\alpha t & \text{if } \lambda=0,\\ \dfrac{\alpha\exp(\lambda t)-\beta}{\lambda} & \text{otherwise,}\end{cases} \tag{12}$$

and

$$p(t)=1/q(t), \tag{13}$$

we can rewrite Eq. (11) as

$$G(s,t)=\left[\frac{p(t)}{1-(1-p(t))s}\right]^\rho, \tag{14}$$

which is the p.g.f. of

$$\text{Negative-Binomial}(\rho,p(t)). \tag{15}$$

Therefore, the p.d.f. of $X(t)$ is

$$P(X(t)=k)=\binom{k+\rho-1}{k}(1-p(t))^k p(t)^\rho. \tag{16}$$

Next, we want to obtain a large-time approximation of $X(t)$, which varies depending on the value of $\lambda$. If $\lambda<0$, then

$$q(t)\to-\frac{\beta}{\lambda} \qquad \text{and} \qquad p(t)\to-\frac{\lambda}{\beta} \qquad \text{as} \qquad t\to\infty. \tag{17}$$

Hence,

$$\lim_{t\to\infty}X(t)=Y\sim\text{Negative-Binomial}\left(\rho,-\frac{\lambda}{\beta}\right) \tag{18}$$

as required. If $\lambda\geq 0$, then

$$q(t)\to\infty \qquad \text{and} \qquad p(t)\to 0 \qquad \text{as} \qquad t\to\infty, \tag{19}$$

which implies

$$\mathbb{E}[X(t)]=\frac{\rho(1-p(t))}{p(t)}\to\infty \qquad \text{as} \qquad t\to\infty. \tag{20}$$

This means that in the regime of large $t$, $P(X(t)=k)$ is only non-trivial for large $k$. Hence, it is sufficient to look at the approximation of $P(X(t)=k)$ as $k\to\infty$. Rewriting Eq. (16) gives

$$P(X(t)=k)=\frac{(k+\rho-1)!}{k!(\rho-1)!}(1-p(t))^k p(t)^\rho=\frac{p(t)^\rho}{\Gamma(\rho)}\frac{(k+\rho-1)!}{k!}(1-p(t))^k. \tag{21}$$

By Stirling's approximation for large $k$,

$$\frac{(k+\rho-1)!}{k!} \sim \frac{\sqrt{2\pi(k+\rho-1)}(k+\rho-1)^{k+\rho-1}/e^{k+\rho-1}}{\sqrt{2\pi k}k^k/e^k}, \tag{22}$$

where

$$\begin{aligned}
\frac{\sqrt{2\pi(k+\rho-1)}}{\sqrt{2\pi k}} &= \sqrt{\frac{k+\rho-1}{k}} \to 1, \\
\frac{(k+\rho-1)^{k+\rho-1}}{k^k} &= \frac{(k+\rho-1)^k(k+\rho-1)^{\rho-1}}{k^k} \to \left(1+\frac{\rho-1}{k}\right)^k k^{\rho-1}.
\end{aligned} \tag{23}$$

Therefore,

$$\begin{aligned}
P(X(t)=k) &\sim \frac{p(t)^\rho}{\Gamma(\rho)}\frac{k^{\rho-1}}{e^{\rho-1}}\left(1+\frac{\rho-1}{k}\right)^k\left(1+\frac{-p(t)k}{k}\right)^k \\
&\sim \frac{p(t)^\rho}{\Gamma(\rho)}\frac{k^{\rho-1}}{e^{\rho-1}}e^{\rho-1}e^{-p(t)k} \\
&= \frac{p(t)^\rho}{\Gamma(\rho)}k^{\rho-1}e^{-p(t)k},
\end{aligned} \tag{24}$$

which is the p.d.f. of the gamma distribution with shape $\rho$ and rate $p(t)$. Since the c.d.f. of $X(t)$ is

$$P(X(t) \le k) = \frac{\gamma(\rho, p(t)k)}{\Gamma(\rho)}, \tag{25}$$

where $\gamma$ is the lower incomplete gamma function, we then have

$$P(X(t) \le q(t)k) = P(X(t) \le p(t)^{-1}k) = \frac{\gamma(\rho, k)}{\Gamma(\rho)}. \tag{26}$$

Hence, as $t \to \infty$,

$$X(t) \to q(t)Z \qquad \text{where} \qquad Z \sim \text{Gamma}(\rho, 1). \tag{27}$$

Furthermore, if $\lambda = 0$, $q(t)$ is dominated by the term $\alpha t$ for large $t$, which gives

$$\lim_{t\to\infty} t^{-1}X(t) = Y = \alpha Z \sim \text{Gamma}\left(\rho, \frac{1}{\alpha}\right). \tag{28}$$

Likewise, if $\lambda > 0$, $q(t)$ is dominated by $\frac{\alpha}{\lambda}\exp(\lambda t)$ for large $t$, and

$$\lim_{t\to\infty} \exp(-\lambda t)X(t) = Y = \frac{\alpha}{\lambda}Z \sim \text{Gamma}\left(\rho, \frac{\lambda}{\alpha}\right) \tag{29}$$

as required. □

Next, we adapt the theories developed in [1] to derive the distributions of the subclonal populations in tumor mutation trees. Note that we focus on the approximate model in the limit of large times and small mutation rates as defined in [1], thereby not introducing different notations to distinguish between the exact and the approximate model. To be consistent with the notations in [1], we define the following quantities:

- Net growth rate of cells in subclone $v$:

$$\lambda_v = \alpha_v - \beta_v$$

- Running-max net growth rate along the trajectory of $v$:

$$\delta_v = \max\{\lambda'_{v'} : v' \in \text{anc}(v) \cup \{v\}\}$$

- Number of times the running-max net growth rate has been attained along the trajectory of $v$:

$$r_v = \#\{v' : v' \in \mathrm{anc}(v), \lambda_{v'} = \delta_v\}$$

- Shape of the population distribution of subclone $v$:

$$\rho_v = \nu_v / \alpha_v$$

- Scale of the population distribution of subclone $v$:

$$\phi_v = \begin{cases} -\beta_v/\lambda_v & \text{if } \lambda_v < 0 \\ \alpha_v & \text{if } \lambda_v = 0 \\ \alpha_v/\lambda_v & \text{if } \lambda_v > 0 \end{cases}$$

- Growth ratio of subclone $v$:

$$\gamma_v = \delta_{\mathrm{pa}(v)}/\delta_v$$

All key notations used in this article are summarized in Table S1.

**Theorem 1.** *Let $\mathscr{T}$ be a tumor mutation tree. For all subclones $v \in \mathscr{T}, v \neq v_0$, there exists a random variable $\tilde{C}_v$ defined on either $\{0, 1, 2, \dots\}$ or $(0, \infty)$ such that*

$$\lim_{t\to\infty} t^{-(r_v-1)} \exp(-\delta_v t) C_v(t) = \tilde{C}_v \tag{30}$$

*almost surely.*

*Proof.* We first consider the base case where the subclone $v$ follows directly after the root, *i.e.* $v \in \mathscr{T}, v \neq v_0$ and $\mathrm{pa}(v) = v_0$. By our model definition, the reactions associated with $v$ are

$$\begin{aligned} C_{v_0} &\xrightarrow{\nu_v} C_{v_0} + C_v \\ C_v &\xrightarrow{\nu_{v'}} C_v + C_{v'} \qquad \text{for all } v' \in \mathrm{ch}(v) \\ C_v &\xrightarrow{\alpha_v} C_v + C_v \\ C_v &\xrightarrow{\beta_v} \emptyset \end{aligned} \tag{31}$$

Here, the wild-type compartment has a static population size, $C_{v_0}(t) = C_0$ for all $t \geq 0$, since the first line of reaction does not increase or decrease the number of wild-type cells. This implies that $\lambda_{v_0} = 0$. Also, the second line of reactions correspond to mutations into the children subclones of $v$ and have no influence on the master equation of $C_v(t)$,

$$\begin{aligned} \frac{dP(C_v(t) = k)}{dt} &= [C_0\nu + \alpha(k-1)]P(C_v(t) = k-1) \\ &\quad + \beta(k+1)P(C_v(t) = k+1) \\ &\quad - [C_0\nu + \alpha k + \beta k]P(C_v(t) = k), \end{aligned} \tag{32}$$

which matches Eq. (6) except for a change of mutation rate from $\nu$ to $C_0\nu$. Therefore, we know from Lemma 1 the large-time limiting distributions of $C_v(t)$:

(i) if $\lambda_v < 0$, then $\delta_v = \lambda_{v_0} = 0$, $r_v = 1$, and

$$\lim_{t\to\infty} t^{-(r_v-1)} \exp(-\delta_v t) C_v(t) = \lim_{t\to\infty} C_v(t) = \tilde{C}_v \sim \text{Negative-Binomial}\left(C_0\rho_v, \phi_v^{-1}\right), \tag{33}$$

(ii) if $\lambda_v = 0$, then $\delta_v = \lambda_{v_0} = \lambda_v = 0$, $r_v = 2$, and

$$\lim_{t\to\infty} t^{-(r_v-1)} \exp(-\delta_v t) C_v(t) = \lim_{t\to\infty} t^{-1} C_v(t) = \tilde{C}_v \sim \text{Gamma}\left(C_0\rho_v, \phi_v^{-1}\right), \tag{34}$$

(iii) if $\lambda > 0$, then $\delta_v = \lambda_v$, $r_v = 1$, and

$$\lim_{t\to\infty} t^{-(r_v-1)} \exp(-\delta_v t) C_v(t) = \lim_{t\to\infty} \exp(-\lambda_v t) C_v(t) = \tilde{C}_v \sim \text{Gamma}\left(C_0\rho_v, \phi_v^{-1}\right), \tag{35}$$

which establish the base case. Next, we observe that for all subclones $v \in \mathscr{T}, v \neq v_0$, the reaction associated with mutations,

$$C_{\text{pa}(v)} \xrightarrow{\nu_v} C_{\text{pa}(v)} + C_v \tag{36}$$

does not change the size of the parent subclone $C_{\text{pa}(v)}$, meaning that all lineages can be viewed as independent linear multi-type branching processes. Therefore, the inductive step for the subclones further downstream follows directly from Proposition 1 & 2 of [1], which concludes the proof. □

**Theorem 2.** *Let $\mathscr{T}$ be a tumor mutation tree. For all subclones $v \in \mathscr{T}, v \neq v_0$ with $\text{pa}(v) \neq v_0$,*

$$\mathbb{E}\left[\exp(-\theta\tilde{C}_v) \mid \tilde{C}_{\text{pa}(v)} = \tilde{c}_{\text{pa}(v)}\right] = \exp\left(-h_v(\theta)\tilde{c}_{\text{pa}(v)}\right), \tag{37}$$

*where*

$$h_v(\theta) = \begin{cases} \dfrac{\nu_v\theta}{\delta_{\text{pa}(v)} - \lambda_v} & \text{if } \delta_{\text{pa}(v)} > \lambda_v \\ \dfrac{\nu_v\theta}{r_{\text{pa}(v)}} & \text{if } \delta_{\text{pa}(v)} = \lambda_v \\ \dfrac{\nu_v\theta(r_{\text{pa}(v)}-1)!}{\lambda_v^{r_{\text{pa}(v)}}}\Phi\left(-\phi_v\theta, r_{\text{pa}(v)}, 1-\gamma_v\right) & \text{if } \delta_{\text{pa}(v)} < \lambda_v, \end{cases} \tag{38}$$

*with $\Phi(z,s,a) = \sum_{k=0}^{\infty} \frac{z^k}{(k+a)^s}$ being the Lerch transcendent function.*

*Proof.* See Corollary 2 of [1]. □

The following proposition provides large $\theta$ approximations of the function $h_v(\theta)$ for the case $\delta_{\text{pa}(v)} < \lambda_v$, which simplifies the calculation of the Lerch transcendent function.

**Proposition 1.** *Consider the case $\delta_{\text{pa}(v)} < \lambda_v$ of $h_v(\theta)$ in Eq. (38). As $\theta \to \infty$, we have the approximation*

$$h_v(\theta) \approx \begin{cases} 2\rho_v \dfrac{(r_{\text{pa}(v)}-1)!}{\lambda_v^{r_{\text{pa}(v)}-1}} \displaystyle\sum_{k=0}^{\lfloor \frac{r_{\text{pa}(v)}}{2} \rfloor} \eta(2k) \dfrac{(\log(\theta\phi_v))^{r_{\text{pa}(v)}-2k}}{(r_{\text{pa}(v)}-2k)!} & \text{if } \gamma_v = 0 \\ \rho_v\phi_v^{\gamma_v} \dfrac{\pi}{\sin(\pi\gamma_v)} \dfrac{\log(\theta\phi_v)^{r_{\text{pa}(v)}-1}}{\lambda_v^{r_{\text{pa}(v)}-1}} \theta^{\gamma_v} & \text{if } \gamma_v > 0 \end{cases} \tag{39}$$

*where $\eta(s) = \sum_{n=1}^{\infty} \frac{(-1)^{n-1}}{n^s}$ is the Dirichlet eta function.*

*Proof.* Suppose $\gamma_v = 0$. Given the property

$$\text{Li}_s(z) = z\Phi(z,s,1), \tag{40}$$

we can rewrite $h_v(\theta)$ as

$$h_v(\theta) = \frac{\nu_v\theta(r_{\text{pa}(v)}-1)!}{\lambda_v^{r_{\text{pa}(v)}}}\left(-\frac{\lambda_v}{\alpha_v\theta}\right)\text{Li}_{r_{\text{pa}(v)}}(-\phi_v\theta) = -\rho_v\frac{(r_{\text{pa}(v)}-1)!}{\lambda_v^{r_{\text{pa}(v)}-1}}\text{Li}_{r_{\text{pa}(v)}}(-\phi_v\theta). \tag{41}$$

By definition, $r_{\text{pa}(v)}$ is a positive integer and $\phi_v > 0$. As $\theta \to \infty$, it follows from eq.(11.1) of [2] that

$$\text{Li}_{r_{\text{pa}(v)}}(-\phi_v\theta) \approx -2\sum_{k=0}^{\lfloor \frac{r_{\text{pa}(v)}}{2} \rfloor} \eta(2k)\frac{(\log(\theta\phi_v))^{r_{\text{pa}(v)}-2k}}{(r_{\text{pa}(v)}-2k)!}, \tag{42}$$

where $\eta(s)=\sum_{n=1}^{\infty}\frac{(-1)^{n-1}}{n^s}$ is the Dirichlet eta function. Hence,

$$h_v(\theta)\approx 2\rho_v\frac{(r_{\mathrm{pa}(v)}-1)!}{\lambda_v^{r_{\mathrm{pa}(v)}-1}}\sum_{k=0}^{\lfloor\frac{r_{\mathrm{pa}(v)}}{2}\rfloor}\eta(2k)\frac{(\log(\theta\phi_v))^{r_{\mathrm{pa}(v)}-2k}}{(r_{\mathrm{pa}(v)}-2k)!}. \tag{43}$$

For the case of $\gamma_v>0$, we apply Lemma 3 of [1] to get

$$\Phi\left(-\phi_v\theta,r_{\mathrm{pa}(v)},1-\gamma_v\right)\approx\frac{\pi}{\sin\left(\pi\gamma_v\right)}\theta^{\gamma_v-1}\phi_v^{\gamma_v-1}\frac{\log(\theta\phi_v)^{r_{\mathrm{pa}(v)}-1}}{(r_{\mathrm{pa}(v)}-1)!} \tag{44}$$

which then gives

$$\begin{aligned}h_v(\theta)&\approx\frac{\nu_v\theta(r_{\mathrm{pa}(v)}-1)!}{\lambda_v^{r_{\mathrm{pa}(v)}}}\frac{\pi}{\sin\left(\pi\gamma_v\right)}\theta^{\gamma_v-1}\phi_v^{\gamma_v-1}\frac{\log(\theta\phi_v)^{r_{\mathrm{pa}(v)}-1}}{(r_{\mathrm{pa}(v)}-1)!}\\&=\rho_v\phi_v^{\gamma_v}\frac{\pi}{\sin\left(\pi\gamma_v\right)}\frac{\log(\theta\phi_v)^{r_{\mathrm{pa}(v)}-1}}{\lambda_v^{r_{\mathrm{pa}(v)}-1}}\theta^{\gamma_v}.\end{aligned} \tag{45}$$

□

**Corollary 1.** *Let $\mathscr{T}$ be a tumor mutation tree. For all subclones $v\in\mathscr{T},v\neq v_0$,*

$$\mathbb{E}\left[\exp(-\theta\tilde{C}_v)\right]=\begin{cases}\left[\phi_{\mathrm{mrca}(v)}+(1-\phi_{\mathrm{mrca}(v)})\exp(-g_v(\theta))\right]^{-C_0\rho_{\mathrm{mrca}(v)}} & \text{if } \lambda_{\mathrm{mrca}(v)}<0\\ \left(1+\phi_{\mathrm{mrca}(v)}g_v(\theta)\right)^{-C_0\rho_{\mathrm{mrca}(v)}} & \text{if } \lambda_{\mathrm{mrca}(v)}\geq 0\end{cases} \tag{46}$$

*where $g_v(\theta)$ satisfies the following recursion: if* $\mathrm{pa}(v)=v_0$*, then $g_v(\theta)=\theta$; otherwise,*

$$g_v(\theta)=\left(g_{\mathrm{pa}(v)}\circ h_v\right)(\theta) \tag{47}$$

*with $h_v(\theta)$ defined in Theorem 2.*

*Proof.* We will prove this by induction. If $\mathrm{pa}(v)=v_0$, then $\mathrm{mrca}(v)=v$ and by Theorem 1,

$$\tilde{C}_v\sim\begin{cases}\text{Negative-Binomial}\left(C_0\rho_v,\phi_v^{-1}\right) & \text{if } \lambda_v<0,\\ \text{Gamma}\left(C_0\rho_v,\phi_v^{-1}\right) & \text{if } \lambda_v\geq 0.\end{cases} \tag{48}$$

It follows that

$$\mathbb{E}\left[\exp(-\theta\tilde{C}_v)\right]=\begin{cases}\left[\phi_v+(1-\phi_v)\exp(-\theta)\right]^{-C_0\rho_v} & \text{if } \lambda_v<0,\\ (1+\phi_v\theta)^{-C_0\rho_v} & \text{if } \lambda_v\geq 0,\end{cases} \tag{49}$$

which proves the base case. Now suppose Eq. (46) is true for any given $v\in\mathscr{T},v\neq v_0$. Let $v'\in\mathrm{ch}(v)$, then $\mathrm{pa}(v')=v$, $\mathrm{mrca}(v')=\mathrm{mrca}(v)$. By Theorem 2, if $\rho_{\mathrm{mrca}(v)}=\rho_{\mathrm{mrca}(v')}\geq 0$, then

$$\begin{aligned}\mathbb{E}\left[\exp(-\theta\tilde{C}_{v'})\right]&=\mathbb{E}\left[\exp\left(-h_{v'}(\theta)\tilde{C}_v\right)\right]\\&=\left(1+\phi_{\mathrm{mrca}(v)}g_v(h_{v'}(\theta))\right)^{-C_0\rho_{\mathrm{mrca}(v)}}\\&=\left(1+\phi_{\mathrm{mrca}(v')}g_{v'}(\theta)\right)^{-C_0\rho_{\mathrm{mrca}(v')}}.\end{aligned} \tag{50}$$

Similarly, if $\rho_{\mathrm{mrca}(v)}=\rho_{\mathrm{mrca}(v')}<0$, then

$$\mathbb{E}\left[\exp(-\theta\tilde{C}_{v'})\right]=\left[\phi_{\mathrm{mrca}(v')}+(1-\phi_{\mathrm{mrca}(v')})\exp(-g_{v'}(\theta))\right]^{-C_0\rho_{\mathrm{mrca}(v')}}, \tag{51}$$

which concludes the proof. □

#### B.2 Distribution of sampling times

**Theorem 3.** *Let $\mathscr{T}$ be a tumor mutation tree, $T_s$ be the sampling time of $\mathscr{T}$, and $C_{sampling} > 0$ be a constant scaling factor of the tumor size at sampling time. Conditioned on the subclonal population sizes $C_v(t), v \in \mathscr{T}, v \neq v_0$ for some time $t \gg 0$, the large-time limiting probability distribution of $T_s$ is*

$$P(T_s > t \mid C_v(t), v \in \mathscr{T}, v \neq v_0) \approx \prod_{v \in \mathscr{T}, v \neq v_0} \exp\left[-\tilde{q}_v(t)\tilde{C}_v\right] \tag{52}$$

*where*

$$\begin{aligned} \tilde{C}_v &= t^{-(r_v-1)} \exp(-\delta_v t) C_v(t), \\ \tilde{q}_v(t) &= \frac{1}{C_{sampling}} \int_0^t s^{r_v-1} \exp(\delta_v s) ds. \end{aligned} \tag{53}$$

*The large-time limiting marginal probability distribution of $T_s$ is*

$$P(T_s > t) \approx \prod_{v \in \text{ch}(v_0)} \left((1 + \phi_v \tilde{g}_v(t))^{-C_0\rho_v}\right)^{\mathbb{1}\{\lambda_v \geq 0\}} \left([\phi_v + (1-\phi_v)\exp(-\tilde{g}_v(t))]^{-C_0\rho_v}\right)^{\mathbb{1}\{\lambda_v < 0\}}, \tag{54}$$

*where*

$$\tilde{g}_v(t) = \tilde{q}_v(t) + \sum_{v' \in \text{ch}(v)} (h_{v'} \circ \tilde{g}_{v'})(t) \tag{55}$$

*with $h_v$ defined in Theorem 2.*

*Proof.* Recall that the sampling event $S$ occurs at rate $\frac{C_{\text{tumor}}(t)}{C_{\text{sampling}}}$, where $C_{\text{tumor}}(t) = \sum_{v \in \mathscr{T}, v \neq v_0} C_v(t)$ is the total population size of the tree $\mathscr{T}$. Conditioned on the subclonal population sizes $C_v(t), v \in \mathscr{T}, v \neq v_0$, the sampling time $T_s$, defined as the first arrival time of $S$, comes from a Poisson process on $[0, \infty)$ with intensity

$$\begin{aligned} \int_0^t \frac{C_{\text{tumor}}(s)}{C_{\text{sampling}}} ds &= \frac{1}{C_{\text{sampling}}} \int_0^t \sum_{v \in \mathscr{T}, v \neq v_0} C_v(s) ds \\ &\approx \frac{1}{C_{\text{sampling}}} \int_0^t \sum_{v \in \mathscr{T}, v \neq v_0} s^{r_v-1} \exp(\delta_v s) \tilde{C}_v ds \qquad \text{by Theorem 1} \\ &= \sum_{v \in \mathscr{T}, v \neq v_0} \left(\frac{1}{C_{\text{sampling}}} \int_0^t s^{r_v-1} \exp(\delta_v s) ds\right) \tilde{C}_v. \end{aligned} \tag{56}$$

It follows that

$$P(T_s > t \mid C_v(t), v \in \mathscr{T}, v \neq v_0) \approx \exp\left(-\sum_{v \in \mathscr{T}, v \neq v_0} \tilde{q}_v(t)\tilde{C}_v\right) = \prod_{v \in \mathscr{T}, v \neq v_0} \exp\left[-\tilde{q}_v(t)\tilde{C}_v\right], \tag{57}$$

where

$$\tilde{q}_v(t) = \frac{1}{C_{\text{sampling}}} \int_0^t s^{r_v-1} \exp(\delta_v s) ds. \tag{58}$$

Let $\tilde{C}_{\mathscr{T}} := \{\tilde{C}_v : v \in \mathscr{T}, v \neq v_0\}$, $d := |V(\mathscr{T}) \setminus \{v_0\}|$, and $D := [0, \infty)^d$. Then,

$$\begin{aligned} P(T_s > t) &\approx \mathbb{E}\left[\prod_{v \in \mathscr{T}, v \neq v_0} \exp\left[-\tilde{q}_v(t)\tilde{C}_v\right]\right] \\ &= \int_D \left(\underbrace{P(\tilde{C}_v = \tilde{c}_v, v \in \mathscr{T}, v \neq v_0)}_{=\prod_{v \in \mathscr{T}, v \neq v_0} P\left(\tilde{C}_v = \tilde{c}_v | \tilde{C}_{\text{pa}(v)} = \tilde{c}_{\text{pa}(v)}\right)} \prod_{v \in \mathscr{T}, v \neq v_0} \exp\left[-\tilde{q}_v(t)\tilde{c}_v\right]\right) d\tilde{c}_{\mathscr{T}}, \end{aligned} \tag{59}$$

where in the limit of small mutations rates the joint probability distribution can be decomposed as a product of conditional distributions of each node given its parent, and $P(\tilde{C}_v = \tilde{c}_v \mid \tilde{C}_{\mathrm{pa}(v)} = \tilde{c}_{\mathrm{pa}(v)}) = P(\tilde{C}_v = \tilde{c}_v)$ if $\mathrm{pa}(v) = v_0$. By grouping related integrals, we can rewrite the above equation as

$$\begin{aligned} P(T_s > t) &\approx \int_D \left( \prod_{v \in \mathscr{T}, v \neq v_0} P\left(\tilde{C}_v = \tilde{c}_v \mid \tilde{C}_{\mathrm{pa}(v)} = \tilde{c}_{\mathrm{pa}(v)}\right) \exp\left[-\tilde{q}_v(t)\tilde{c}_v\right] \right) d\tilde{c}_{\mathscr{T}} \\ &= \prod_{v \in \mathrm{ch}(v_0)} I_v, \end{aligned} \tag{60}$$

where $I_v$ satisfies the recursion

$$I_v = \begin{cases} \displaystyle\int_0^\infty P\left(\tilde{C}_v = \tilde{c}_v \mid \tilde{C}_{\mathrm{pa}(v)} = \tilde{c}_{\mathrm{pa}(v)}\right) \exp\left[-\tilde{q}_v(t)\tilde{c}_v\right] \left( \prod_{v' : v' \in \mathrm{ch}(v)} I_{v'} \right) d\tilde{c}_v & \text{if } \mathrm{ch}(v) \neq \emptyset, \\ \displaystyle\int_0^\infty P\left(\tilde{C}_v = \tilde{c}_v \mid \tilde{C}_{\mathrm{pa}(v)} = \tilde{c}_{\mathrm{pa}(v)}\right) \exp\left[-\tilde{q}_v(t)\tilde{c}_v\right] d\tilde{c}_v & \text{otherwise.} \end{cases} \tag{61}$$

Let us define

$$\tilde{g}_v(t) = \tilde{q}_v(t) + \sum_{v' \in \mathrm{ch}(v)} (h_{v'} \circ \tilde{g}_{v'})(t). \tag{62}$$

If $v$ is a leaf of $\mathscr{T}$, *i.e.* $\mathrm{ch}(v) = \emptyset$, then by Theorem 2,

$$\begin{aligned} I_v &= \mathbb{E}\left[\exp(-\tilde{q}_v(t)\tilde{C}_v) \mid \tilde{C}_{\mathrm{pa}(v)} = \tilde{c}_{\mathrm{pa}(v)}\right] \\ &= \exp\left(-h_v(\tilde{q}_v(t))\tilde{c}_{\mathrm{pa}(v)}\right) \\ &= \exp\left(-(h_v \circ \tilde{g}_v)(t)\tilde{c}_{\mathrm{pa}(v)}\right). \end{aligned} \tag{63}$$

Therefore, in the case of $\mathrm{ch}(v) \neq \emptyset$, we have

$$\begin{aligned} I_v &= \int_0^\infty P\left(\tilde{C}_v = \tilde{c}_v \mid \tilde{C}_{\mathrm{pa}(v)} = \tilde{c}_{\mathrm{pa}(v)}\right) \exp\left[-\tilde{q}_v(t)\tilde{c}_v\right] \left( \prod_{v' : v' \in \mathrm{ch}(v)} \exp\left(-(h_{v'} \circ \tilde{g}_{v'})(t)\tilde{c}_v\right) \right) d\tilde{c}_v \\ &= \int_0^\infty P\left(\tilde{C}_v = \tilde{c}_v \mid \tilde{C}_{\mathrm{pa}(v)} = \tilde{c}_{\mathrm{pa}(v)}\right) \exp\left[ -\left( \tilde{q}_v(t) + \sum_{v' : v' \in \mathrm{ch}(v)} (h_{v'} \circ \tilde{g}_{v'})(t) \right) \tilde{c}_v \right] d\tilde{c}_v \\ &= \int_0^\infty P\left(\tilde{C}_v = \tilde{c}_v \mid \tilde{C}_{\mathrm{pa}(v)} = \tilde{c}_{\mathrm{pa}(v)}\right) \exp\left[-\tilde{g}_v(t)\tilde{c}_v\right] d\tilde{c}_v \\ &= \mathbb{E}\left[\exp(-\tilde{g}_v(t)\tilde{C}_v) \mid \tilde{C}_{\mathrm{pa}(v)} = \tilde{c}_{\mathrm{pa}(v)}\right]. \end{aligned} \tag{64}$$

For all $v \in \mathrm{ch}(v_0)$, by Corollary 1,

$$I_v = \begin{cases} \left[\phi_v + (1 - \phi_v)\exp(-\tilde{g}_v(t))\right]^{-C_0 \rho_v} & \text{if } \lambda_v < 0, \\ \left(1 + \phi_v \tilde{g}_v(t)\right)^{-C_0 \rho_v} & \text{if } \lambda_v \geq 0, \end{cases} \tag{65}$$

which concludes the proof. □

#### B.3 Expected subclone sizes

For any subclone $v$, the expected subclone size given the parent size $c_{\mathrm{pa}(v)}(t)$ at time $t$ is

$$\mathbb{E}\left[C_v(t) \mid C_{\mathrm{pa}(v)}(t) = c_{\mathrm{pa}(v)}(t)\right] \approx t^{r_v - 1} \exp(\delta_v t) \mathbb{E}\left[\tilde{C}_v \mid \tilde{C}_{\mathrm{pa}(v)} = \tilde{c}_{\mathrm{pa}(v)}\right] \tag{66}$$

by Theorem 1. Given the Laplace transform in Theorem 2, the first moment is

$$\begin{aligned}\tilde{\mu}_v := \mathbb{E}\left[\tilde{C}_v \mid \tilde{C}_{\mathrm{pa}(v)} = \tilde{c}_{\mathrm{pa}(v)}\right] &= -\frac{d}{d\theta}\mathbb{E}\left[\exp(-\theta\tilde{C}_v) \mid \tilde{C}_{\mathrm{pa}(v)} = \tilde{c}_{\mathrm{pa}(v)}\right]\Big|_{\theta=0} \\ &= -\frac{d}{d\theta}\exp(-h_v(\theta)\tilde{c}_{\mathrm{pa}(v)})\Big|_{\theta=0} \\ &= \tilde{c}_{\mathrm{pa}(v)} h'_v(\theta)\exp(-h_v(\theta)\tilde{c}_{\mathrm{pa}(v)})\big|_{\theta=0}.\end{aligned} \tag{67}$$

From Eq. (38) we have

$$h_v(0) = 0 \tag{68}$$

and

$$h'_v(0) = \begin{cases} \dfrac{\nu_v}{\delta_{\mathrm{pa}(v)} - \lambda_v} & \text{if } \delta_{\mathrm{pa}(v)} > \lambda_v, \\ \dfrac{\nu_v}{r_{\mathrm{pa}(v)}} & \text{if } \delta_{\mathrm{pa}(v)} = \lambda_v, \\ \dfrac{\nu_v(r_{\mathrm{pa}(v)} - 1)!}{(\lambda_v - \delta_{\mathrm{pa}(v)})^{r_{\mathrm{pa}(v)}}} & \text{if } \delta_{\mathrm{pa}(v)} < \lambda_v. \end{cases} \tag{69}$$

If follows that

$$\begin{aligned}&\mathbb{E}\left[C_v(t) \mid C_{\mathrm{pa}(v)}(t) = c_{\mathrm{pa}(v)}(t)\right] \\ &\approx t^{r_v - 1}\exp(\delta_v t)\nu_v \tilde{c}_{\mathrm{pa}(v)} h'_v(0) \\ &= \nu_v c_{\mathrm{pa}(v)}(t) \times \begin{cases} \dfrac{1}{\delta_{\mathrm{pa}(v)} - \lambda_v} & \text{if } \delta_{\mathrm{pa}(v)} > \lambda_v, \\ \dfrac{t}{r_{\mathrm{pa}(v)}} & \text{if } \delta_{\mathrm{pa}(v)} = \lambda_v, \\ \dfrac{t^{r_v - r_{\mathrm{pa}(v)}} e^{(\delta_v - \delta_{\mathrm{pa}(v)})t}(r_{\mathrm{pa}(v)} - 1)!}{(\lambda_v - \delta_{\mathrm{pa}(v)})^{r_{\mathrm{pa}(v)}}} & \text{if } \delta_{\mathrm{pa}(v)} < \lambda_v. \end{cases}\end{aligned} \tag{70}$$

### C Parameter inference

#### C.1 Log-likelihood

Recall that the joint log-likelihood of observing a tumor mutation tree $\mathscr{T}$ and sampling time $t_s$ is given by

$$\begin{aligned}\log p(\mathscr{T}, t_s \mid \Theta) = &\sum_{v \in \mathrm{ch}(v_0)} \log p(c_v(t_s) \mid \Theta) \\ &+ \sum_{v \notin \mathrm{ch}(v_0)} \log p(c_v(t_s) \mid c_{\mathrm{pa}(v)}(t_s), \Theta) \\ &+ \log p(t_s \mid c_v(t_s), v \in \mathscr{T}, \Theta) \\ &+ \log p\left(\mathbf{c}'_{\mathscr{T}}(t_s) \mid \mathbf{c}_{\mathscr{T}}(t_s)\right)\end{aligned} \tag{71}$$

The actual subclone sizes $\mathbf{C}_{\mathscr{T}}(t)$ are unobservable. Assuming that the sample proportions $C'_v(t_s)/C_{\mathrm{seq}}$ accurately represent the true cancer cell fractions $C_v(t_s)/C_{\mathrm{tumor}}(t_s)$ in patients, we multiply the sample proportions by the tumor size $C_{\mathrm{tumor}}(t_s)$ to obtain an estimate of $\mathbf{C}_{\mathscr{T}}(t)$. The first summation in Eq. (71) corresponds to the log-likelihood of all nodes directly following the root $v_0$ of $\mathscr{T}$, which can be computed exactly using the Negative-Binomial p.m.f. as given in Eq. (16). The second summation corresponds to the conditional log-likelihood of the remaining nodes given their parents in $\mathscr{T}$. For each node, we approximates the conditional p.m.f. of $c_v(t_s)$ given $c_{\mathrm{pa}(v)}(t_s)$ by numerically inverting the conditional Laplace transform provided in Theorem 2,

$$\begin{aligned}p(c_v(t_s) \mid c_{\mathrm{pa}(v)}(t_s), \Theta) \approx &\mathcal{L}^{-1}\left\{\frac{1}{\theta}\exp\left(-h_v(\theta)\tilde{c}_{\mathrm{pa}(v)}\right)\right\}(\tilde{c}_v^+) \\ &- \left(\mathcal{L}^{-1}\left\{\frac{1}{\theta}\exp\left(-h_v(\theta)\tilde{c}_{\mathrm{pa}(v)}\right)\right\}(\tilde{c}_v)\right)\mathbb{1}\{\tilde{c}_v \neq 0\},\end{aligned} \tag{72}$$

where the cell numbers are converted to their time-independent counterparts according to Theorem 1,

$$\begin{aligned}\tilde{c}_v^+ &= t_s^{-(r_v-1)}\exp(-\delta_v t_s)(c_v(t_s)+1), \\ \tilde{c}_v &= t_s^{-(r_v-1)}\exp(-\delta_v t_s)c_v(t_s), \\ \tilde{c}_{\mathrm{pa}(v)} &= t_s^{-(r_{\mathrm{pa}(v)}-1)}\exp(-\delta_{\mathrm{pa}(v)} t_s)c_{\mathrm{pa}(v)}(t_s).\end{aligned} \tag{73}$$

Here we take the difference in the c.d.f. of the large-time limiting random variable $\tilde{C}_v$ at points $\tilde{c}_v^+$ and $\tilde{c}_v$, since the support of $C_v(t)$ is on the non-negative integers $\{0, 1, 2, \dots\}$, whereas $\tilde{C}_v$ is positive continuous. For the numerical inverse Laplace transform, we use the concentrated matrix exponential method as described in [3] to avoid oscillations and preserve monotonicity in the estimated c.d.f.'s. We note that for the cases when $v$ is equally or less fit than its ancestor subclones ($\lambda_v \leq \delta_{\mathrm{pa}(v)}$), the conditional inverse Laplace transforms in Theorem 2 reduce to Dirac delta functions concentrated on the asymptotic means Eq. (70), which causes difficulties in inference, as any deviations from the means would lead to zero probabilities. We address this issue in Section C.2. We compute the conditional p.d.f. of the sampling time given subclonal population sizes using Theorem 3,

$$\begin{aligned}p(t_s \mid c_v(t_s), v \in \mathscr{T}, \Theta) &= -\frac{d}{dt}P(T_s > t_s \mid c_v(t), v \in \mathscr{T}, v \neq v_0) \\ &\approx \frac{c_{\mathrm{tumor}}(t_s)}{C_{\mathrm{sampling}}}\exp\left(-\sum_{v \in \mathscr{T}, v \neq v_0}\tilde{q}_v(t_s)\tilde{c}_v\right),\end{aligned} \tag{74}$$

which gives

$$\log p(t_s \mid c_v(t_s), v \in \mathscr{T}, \Theta) = \log c_{\mathrm{tumor}}(t_s) - \log C_{\mathrm{sampling}} - \sum_{v \in \mathscr{T}, v \neq v_0}\tilde{q}_v(t_s)\tilde{c}_v. \tag{75}$$

The last term in Eq. (71) is the log density of sampling $\mathbf{c}'_{\mathscr{T}}(t_s)$ without replacement from $\mathbf{c}_{\mathscr{T}}(t_s)$, which is multivariate hypergeometric,

$$\sum_{v\in\mathscr{T}, v\neq v_0} \log\binom{c_v(t_s)}{c'_v(t_s)} - \log\binom{c_{\text{tumor}}(t_s)}{c_{\text{seq}}}. \tag{76}$$

There is no closed-form formula for the marginal log-likelihood of observing a tumor mutation tree $\mathscr{T}$. Hence, if the sampling time is not available, we need to integrate it out,

$$p(\mathscr{T} \mid \Theta) = \int_0^\infty p(\mathscr{T}, t_s \mid \Theta) dt_s. \tag{77}$$

In section Section C.3 we discuss how to handle the unobserved subclones in the trees when calculating the log-likelihood.

#### C.2 Conditional size distributions for the equally and less fit cases

In this section, we approximate the conditional distributions $C_v(t) \mid C_{\text{pa}(v)}(t) = c_{\text{pa}(v)}(t)$ using the mean and the variance. We consider the large-time limit

$$\lim_{t\to\infty} t^{-(r_{\text{pa}(v)}-1)} e^{-\delta_{\text{pa}(v)} t} c_{\text{pa}(v)}(t) = \tilde{c}_{\text{pa}(v)}. \tag{78}$$

Suppose that $(T_i)_{i\in\mathbb{N}}$ comes from a Poisson process on $[0,\infty)$ with intensity $\nu_v c_{\text{pa}(v)}(\cdot)$ and that $(Y_i(t))_{t\geq 0}$, $i \in \mathbb{N}$ are i.i.d. birth-death process initializing from a single cell with birth rate $\alpha_v > 0$, death date $\beta_v > 0$, and net growth rate $\lambda_v = \alpha_v - \beta_v$. Then, an alternative definition of $C_v(t)$ is

$$C_v(t) = \sum_{i:T_i\leq t} Y_i(t - T_i). \tag{79}$$

Conditioned on $C_{\text{pa}(v)}(t) = c_{\text{pa}(v)}(t)$, the variance of $C_v(t)$ is a known result: if $\lambda_v \neq 0$, then

$$\begin{aligned} \text{Var}(C_v(t)) &= \int_0^t \nu_v c_{\text{pa}(v)}(t) \left( \frac{2\alpha_v}{\lambda_v} e^{2\lambda_v(t-s)} - \frac{\alpha_v+\beta_v}{\lambda_v} e^{\lambda_v(t-s)} \right) ds \\ &\approx \nu_v \tilde{c}_{\text{pa}(v)} \int_0^t s^{r_{\text{pa}(v)}-1} e^{\delta_{\text{pa}(v)} s} \left( \frac{2\alpha_v}{\lambda_v} e^{2\lambda_v(t-s)} - \frac{\alpha_v+\beta_v}{\lambda_v} e^{\lambda_v(t-s)} \right) ds; \end{aligned} \tag{80}$$

if $\lambda_v = 0$, then

$$\begin{aligned} \text{Var}(C_v(t)) &= \int_0^t \nu_v c_{\text{pa}(v)}(t) \left(1 + 2\alpha_v(t-s)\right) ds \\ &\approx \nu_v \tilde{c}_{\text{pa}(v)} \int_0^t s^{r_{\text{pa}(v)}-1} e^{\delta_{\text{pa}(v)} s} \left(1 + 2\alpha_v(t-s)\right) ds. \end{aligned} \tag{81}$$

For the case $\lambda_v < \delta_{\text{pa}(v)}$, we have $r_v = r_{\text{pa}(v)}$ and $\delta_v = \delta_{\text{pa}(v)}$. If follows that

$$\begin{aligned} &\tilde{\sigma}_v^2 = \text{Var}\left(\tilde{C}_v\right) = \text{Var}\left(t^{-(r_v-1)} e^{-\delta_v t} C_v(t)\right) \\ &\approx \nu_v \tilde{c}_{\text{pa}(v)} t^{-2(r_v-1)} \times \\ &\begin{cases} \dfrac{2\alpha_v}{\lambda_v} \displaystyle\int_0^t s^{r_v-1} e^{(\delta_v-2\lambda_v)s-2(\delta_v-\lambda_v)t} ds - \dfrac{\alpha_v+\beta_v}{\lambda_v} \displaystyle\int_0^t s^{r_v-1} e^{(\delta_v-\lambda_v)s-(2\delta_v-\lambda_v)t} ds & \text{if } \lambda_v \neq 0, \\ (1+2\alpha_v t) \displaystyle\int_0^t s^{r_v-1} e^{\delta_v s - 2\delta_v t} ds - 2\alpha_v \displaystyle\int_0^t s^{r_v} e^{\delta_v s - 2\delta_v t} ds & \text{if } \lambda_v = 0. \end{cases} \end{aligned} \tag{82}$$

For the case $\lambda_v = \delta_{\text{pa}(v)}$, we have $r_v = r_{\text{pa}(v)} + 1$ and $\lambda_v = \delta_v = \delta_{\text{pa}(v)}$. Then,

$$\begin{aligned} \tilde{\sigma}_v^2 &= \text{Var}\left(\tilde{C}_v\right) = \text{Var}\left(t^{-(r_v-1)} e^{-\delta_v t} C_v(t)\right) \\ &\approx \nu_v \tilde{c}_{\text{pa}(v)} t^{-2(r_v-1)} \times \begin{cases} \dfrac{2\alpha_v}{\delta_v} \displaystyle\int_0^t s^{r_v-2} e^{-\delta_v s} ds - \dfrac{\alpha_v+\beta_v}{\delta_v} e^{-\delta_v t} \dfrac{t^{r_v-1}}{r_v-1} & \text{if } \lambda_v \neq 0, \\ (1+2\alpha_v t) \dfrac{t^{r_v-1}}{r_v-1} - 2\alpha_v \dfrac{t^{r_v}}{r_v} & \text{if } \lambda_v = 0. \end{cases} \end{aligned} \tag{83}$$

Finally, we use log-normal distributions, where the mean and variance are determined using Eq. (67), Eq. (82), and Eq. (83),

$$\mathrm{Lognormal}(\mu, \sigma^2) \tag{84}$$

$$\sigma^2 = \log\left(1 + \frac{\tilde{\sigma}_v^2}{\tilde{\mu}_v^2}\right) \tag{85}$$

$$\mu = \log\left(\tilde{\mu}_v\right) - \frac{1}{2}\log\left(1 + \frac{\tilde{\sigma}_v^2}{\tilde{\mu}_v^2}\right) = \log\left(\tilde{\mu}_v\right) - \frac{1}{2}\sigma^2 \tag{86}$$

to approximate the conditional distributions for the cases when $\lambda_v \leq \delta_{\mathrm{pa}(v)}$. We show in simulations that these distributions fit reasonably well (Figure S9, Figure S10, Figure S11).

#### C.3 Unobserved subclones, augmented trees, and the union tree

To compute the log-likelihood of a tree, it is essential to account for both observed and unobserved subclones. This means considering all subclones defined by the tree expansion rule $R$, even if no cells are attached. However, the number of summands in Eq. (71) can grow super-exponentially in the number of mutations $n$ depending on $R$. For example, if we allow for mutations to appear in parallel branches, the number of mutant subclones is $\mathcal{O}(n!)$ (Figure S1 (a) & (b)). Therefore, calculating the exact log-likelihood over all subclones defined by $R$ can quickly become computationally intractable as $n$ increases. At the same time, restricting the computation to only the observed subclones introduces substantial bias into the results.

To address this issue, we first augment each tree in the cohort based on the tree expansion rule $R$. Specifically, for each observed subclone in the tree, we add a child subclone whenever its genotype is permissible by $R$. Since these newly added subclones are unobserved, we assume they contain no cells. Next, we compute the union tree of all trees in $\mathscr{T} = \{\mathscr{T}^j\}_{j\in N}$. That is, we construct a single tree that includes all unique subclones across all augmented trees. The summations in Eq. (71) then go through all these subclones in the union tree. We illustrate this process in Figure S1 (c) and (d).

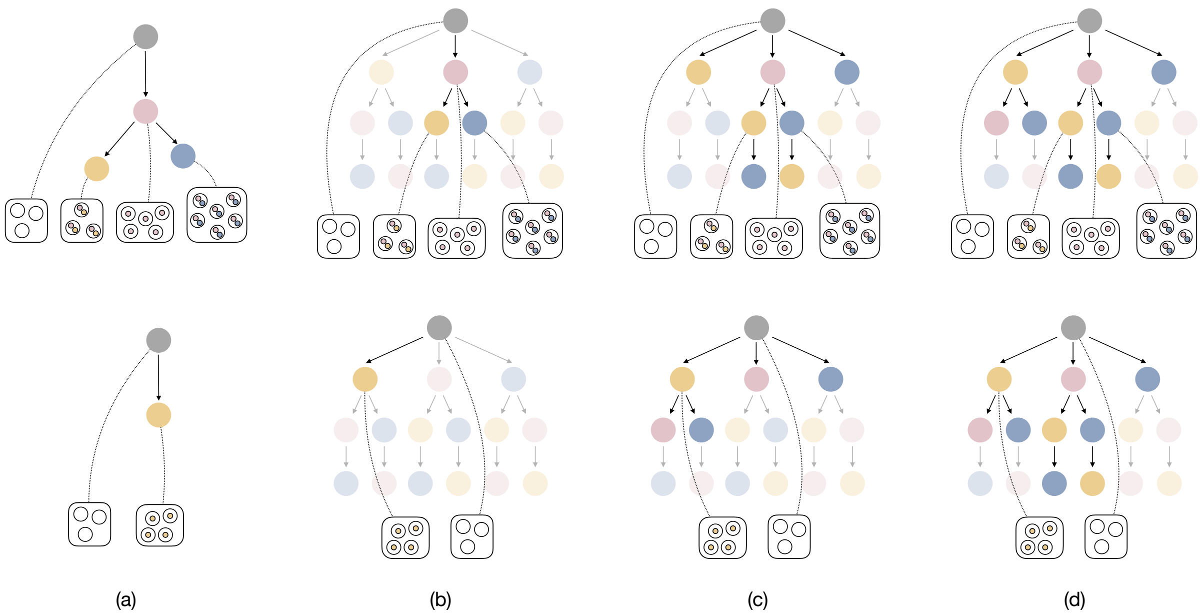

Figure S1: Illustration of converting observed tumor mutation trees for log-likelihood computation. (a) Tumor mutation trees with only observed subclones. (b) Complete trees including unobserved subclones, assuming $n = 3$ mutations and a parallel tree expansion rule. Observed subclones are highlighted, and unobserved ones are more transparent. (c) Augmented trees generated using the parallel expansion rule, with observed and augmented subclones highlighted. (d) Final trees for log-likelihood calculation (Eq. (71)), highlighting the union tree of the augmented trees in (c).

### D Simulation studies

#### D.1 Validation of distributions against stochastic simulations

In this section, we use stochastic simulations to validate our approximations of the marginal distribution functions for subclonal population sizes and sampling times. We consider a linear and a branching tree of three subclones with different parameter settings as summarized in Supplementary Table S2. For setting A through D, F, and G, we simulate under the linear model

$$\begin{aligned}
\emptyset &\xrightarrow{\nu_1} C_1 \\
C_1 &\xrightarrow{\alpha_1} C_1 + C_1 \\
C_1 &\xrightarrow{\beta_1} \emptyset \\
C_1 &\xrightarrow{\nu_2} C_1 + C_2 \\
C_2 &\xrightarrow{\alpha_2} C_2 + C_2 \\
C_2 &\xrightarrow{\beta_2} \emptyset \\
C_2 &\xrightarrow{\nu_3} C_2 + C_3 \\
C_3 &\xrightarrow{\alpha_3} C_3 + C_3 \\
C_3 &\xrightarrow{\beta_3} \emptyset \\
\emptyset &\xrightarrow{\frac{C_1+C_2+C_3}{C_{\text{sampling}}}} S
\end{aligned} \tag{87}$$

For setting E, where a branching tree structure is used, we replace the line $C_2 \xrightarrow{\nu_3} C_2 + C_3$ by $C_1 \xrightarrow{\nu_3} C_1 + C_3$. We perform stochastic simulations using the explicit tau-leap method [4] provided in the `R` package `GillespieSSA2` [5]. For each setting, we choose the tau-leap step size and the end time to be 0.002 and 50 respectively, and we run simulations for 100000 times. In Supplementary Figure S2 through Supplementary Figure S8, we compare the simulation results against our approximations for the marginal distributions of the three cell types at time $t = 20$ and the marginal distribution of the sampling time.

| **Setting** | **Branching** | $C_0$ | $C_{\text{sampling}}$ | $\nu_1$ | $\alpha_1$ | $\beta_1$ | $\nu_2$ | $\alpha_2$ | $\beta_2$ | $\nu_3$ | $\alpha_3$ | $\beta_3$ |
|---|---|---|---|---|---|---|---|---|---|---|---|---|
| A | No | 700 | $10^9$ | 0.001 | 1 | 0.3 | 0.01 | 1 | 1.5 | 0.01 | 1.4 | 0.3 |
| B | No | 700 | $10^9$ | 0.001 | 1 | 0.3 | 0.01 | 1.4 | 0.3 | 0.01 | 1 | 1.5 |
| C | No | 700 | $10^9$ | 0.001 | 1 | 0.3 | 0.01 | 1 | 0.3 | 0.01 | 1 | 0.3 |
| D | No | 700 | $10^9$ | 0.001 | 1 | 0.3 | 0.001 | 1.4 | 0.3 | 0.001 | 1 | 1.5 |
| E | Yes | 700 | $10^9$ | 0.001 | 1 | 0.3 | 0.01 | 1 | 1.5 | 0.001 | 1.4 | 0.3 |
| F | No | 700 | $10^9$ | 0.001 | 0.3 | 0.3 | 0.01 | 1.4 | 0.3 | 0.01 | 1.5 | 0.3 |
| G | No | 700 | $10^9$ | 0.001 | 0.7 | 1.2 | 0.01 | 1.3 | 0.3 | 0.01 | 1.5 | 0.3 |

Table S2: A summary of parameter settings in stochastic simulations.

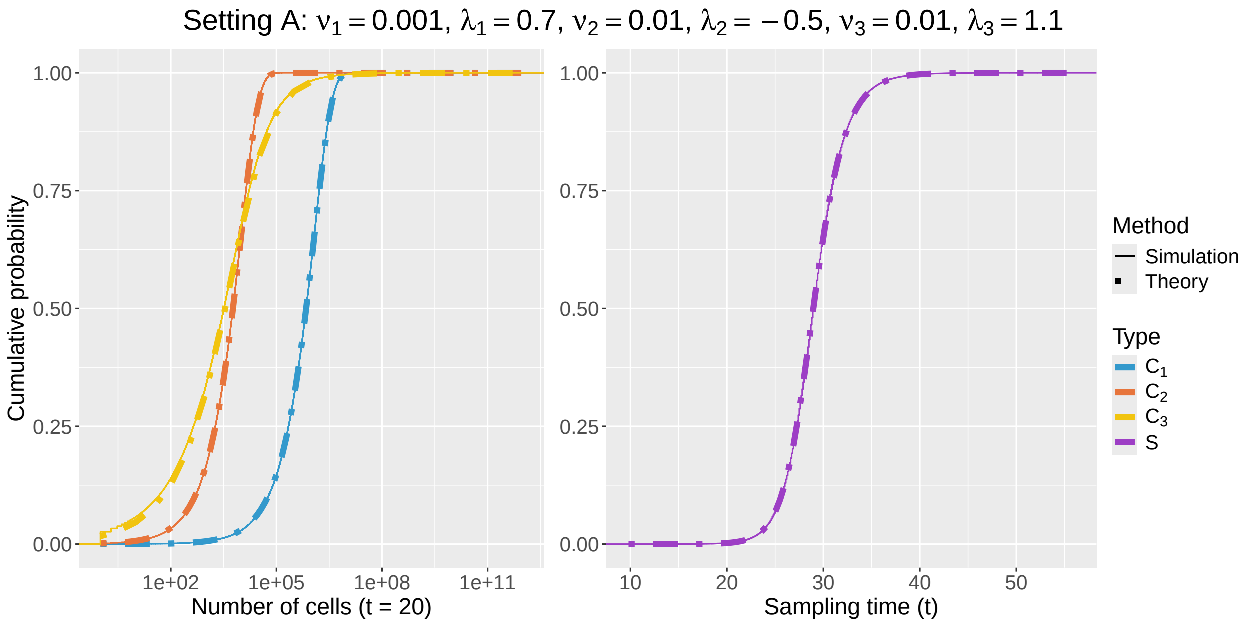

Figure S2: Comparison between the empirical marginal cumulative distributions of subclonal population sizes (left) and sampling time (right) obtained using stochastic simulations (solid lines) and from theoretical approximations (dash lines) under setting A.

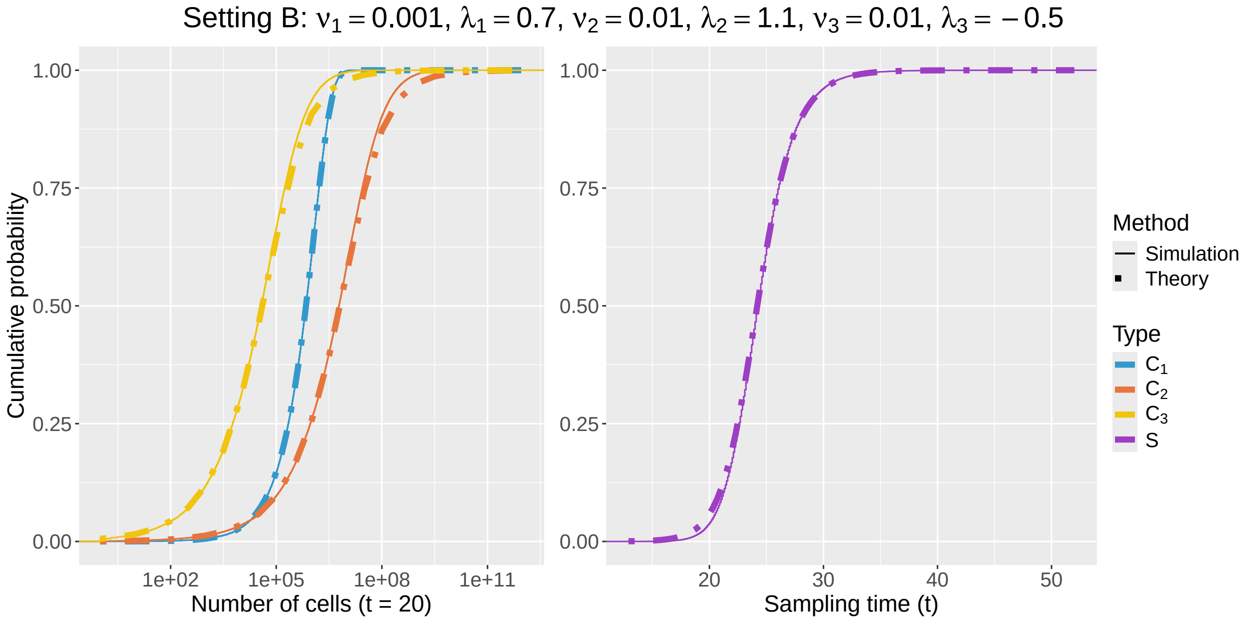

Figure S3: Comparison between the empirical marginal cumulative distributions of subclonal population sizes (left) and sampling time (right) obtained using stochastic simulations (solid lines) and from theoretical approximations (dash lines) under setting B.

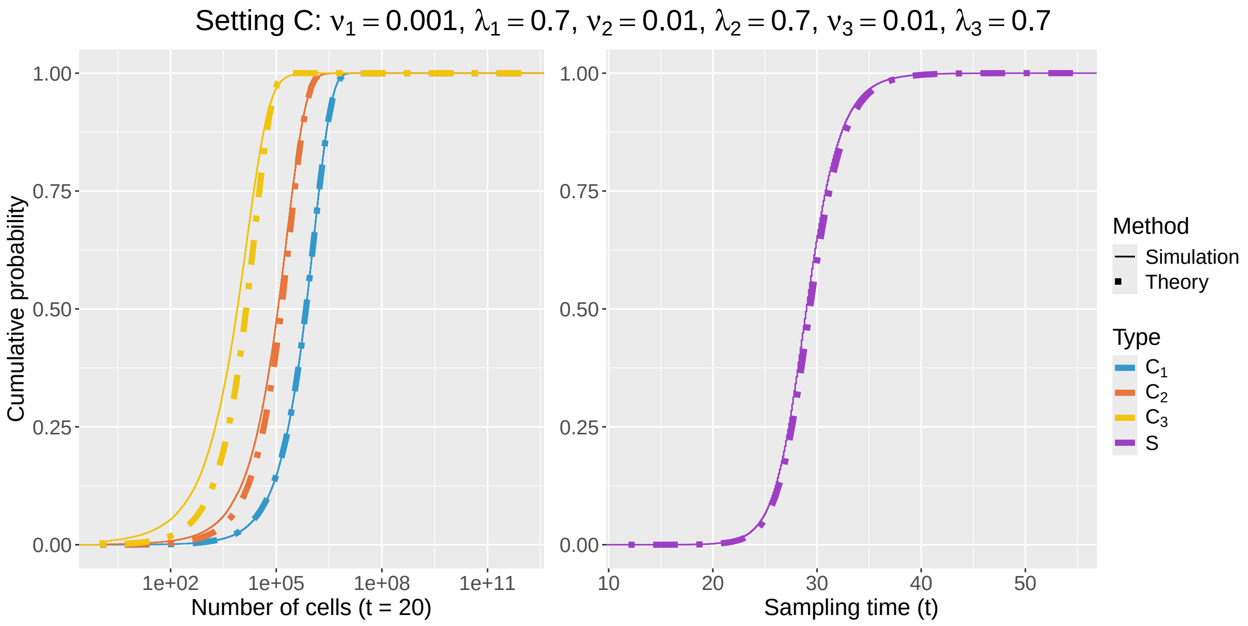

Figure S4: Comparison between the empirical marginal cumulative distributions of subclonal population sizes (left) and sampling time (right) obtained using stochastic simulations (solid lines) and from theoretical approximations (dash lines) under setting C.

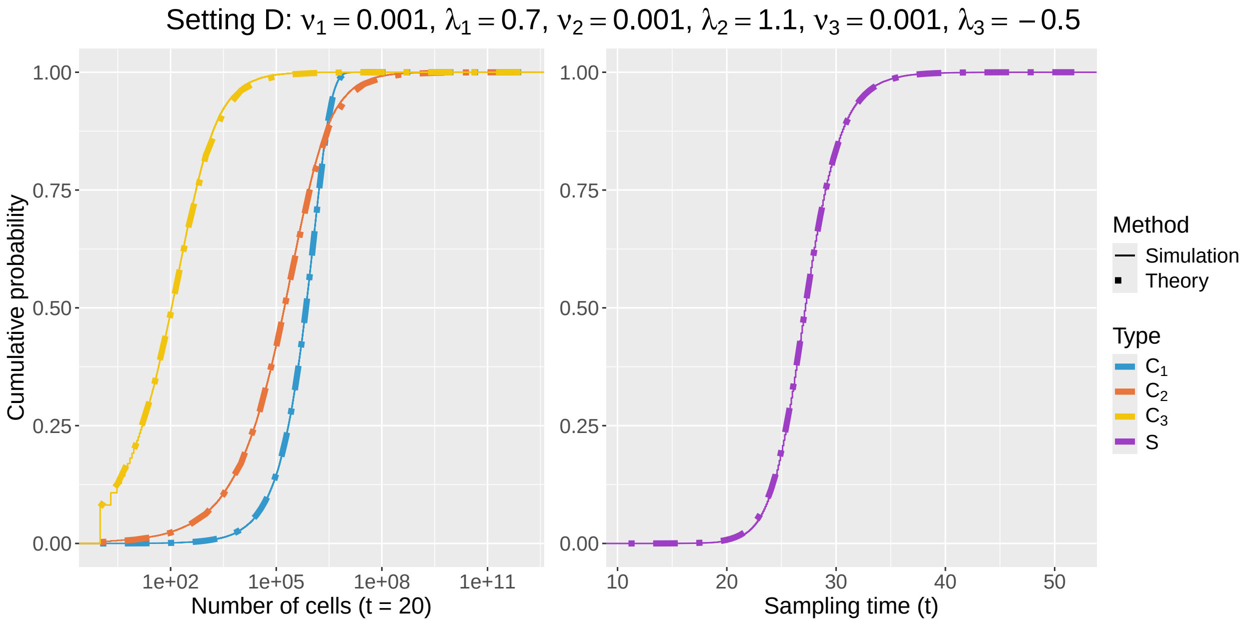

Figure S5: Comparison between the empirical marginal cumulative distributions of subclonal population sizes (left) and sampling time (right) obtained using stochastic simulations (solid lines) and from theoretical approximations (dash lines) under setting D.

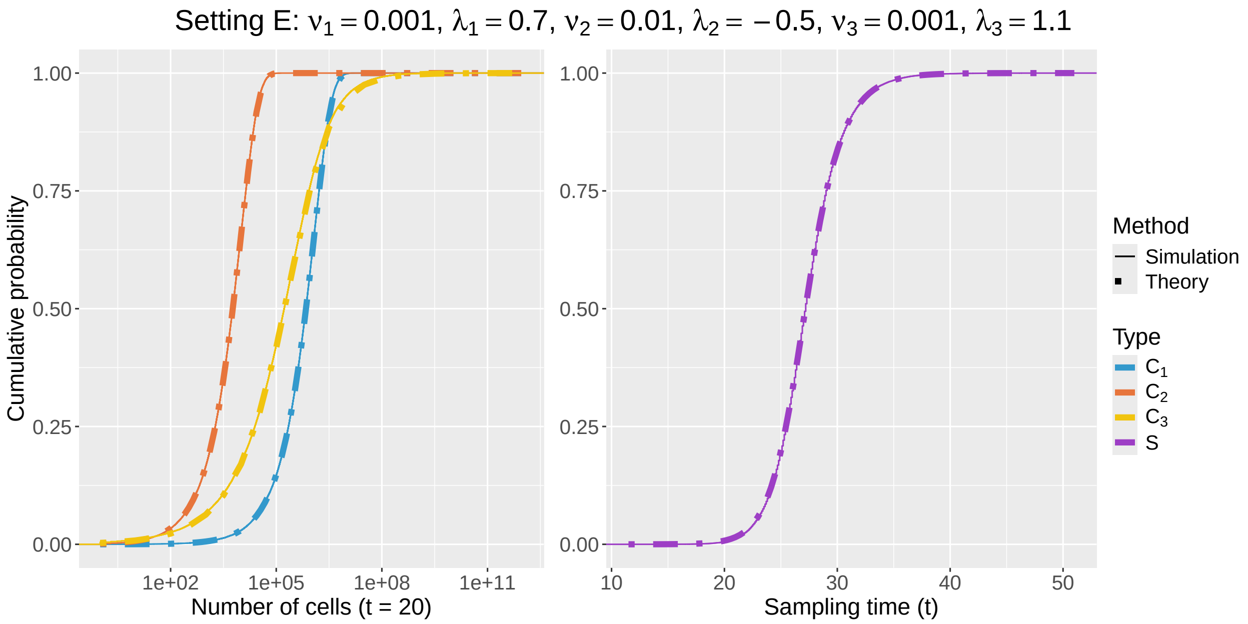

Figure S6: Comparison between the empirical marginal cumulative distributions of subclonal population sizes (left) and sampling time (right) obtained using stochastic simulations (solid lines) and from theoretical approximations (dash lines) under setting E.

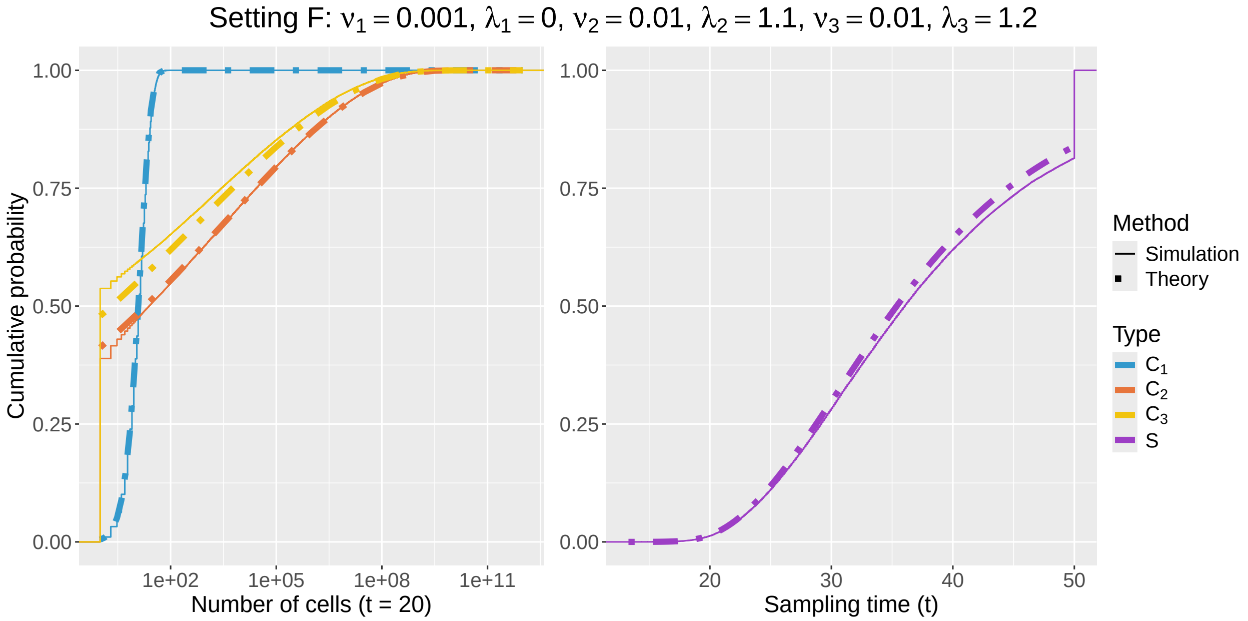

Figure S7: Comparison between the empirical marginal cumulative distributions of subclonal population sizes (left) and sampling time (right) obtained using stochastic simulations (solid lines) and from theoretical approximations (dash lines) under setting F.

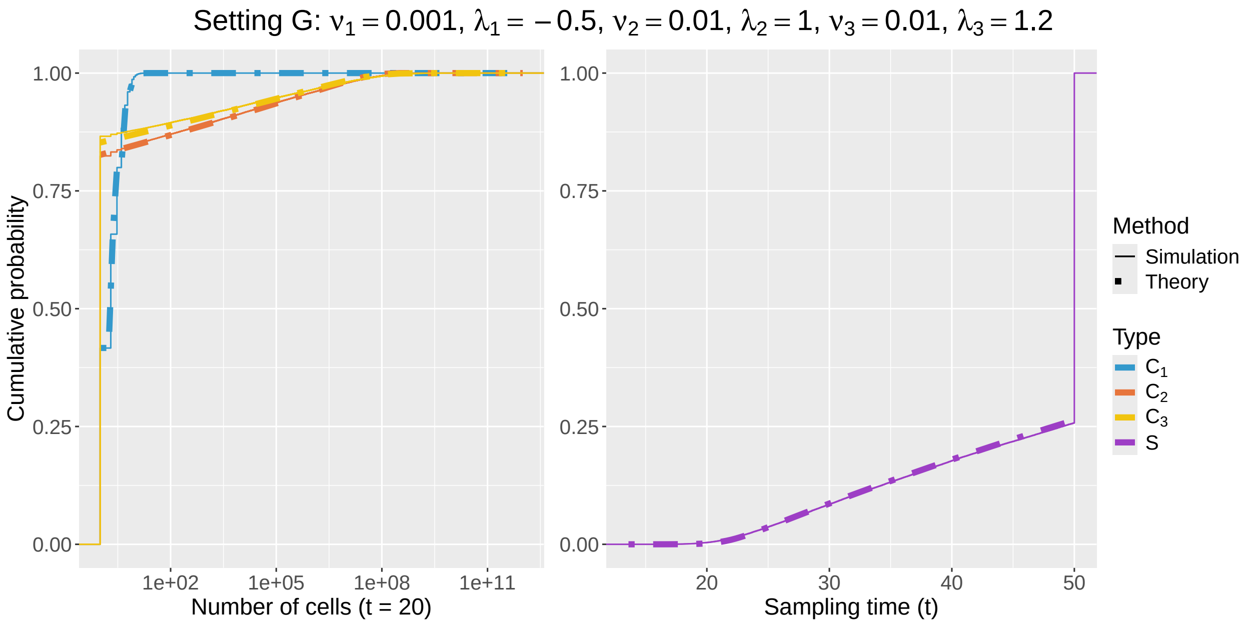

Figure S8: Comparison between the empirical marginal cumulative distributions of subclonal population sizes (left) and sampling time (right) obtained using stochastic simulations (solid lines) and from theoretical approximations (dash lines) under setting G.

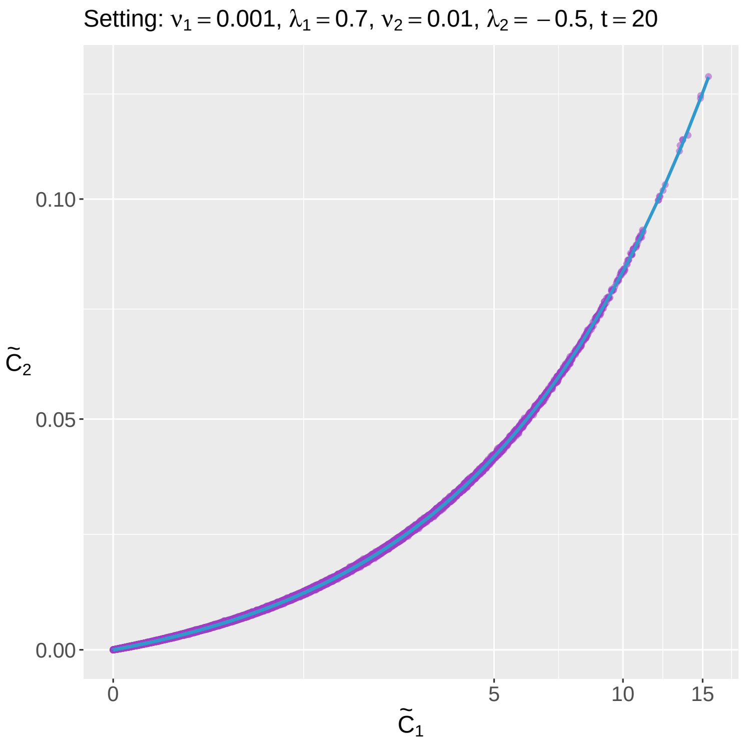

Figure S9: Comparison between the simulated cell numbers and the approximated log-normal distribution for the case $\lambda_1 = 0.7 > -0.5 = \lambda_2$. The cell numbers have been converted to their time-independent counterparts based on Theorem 1. The purple dots are the simulated values. The blue line is the theoretical mean given in Eq. (67). The blue region corresponds to the interval between the 95% and 5% quantiles of the log-normal distribution defined in Eq. (84).

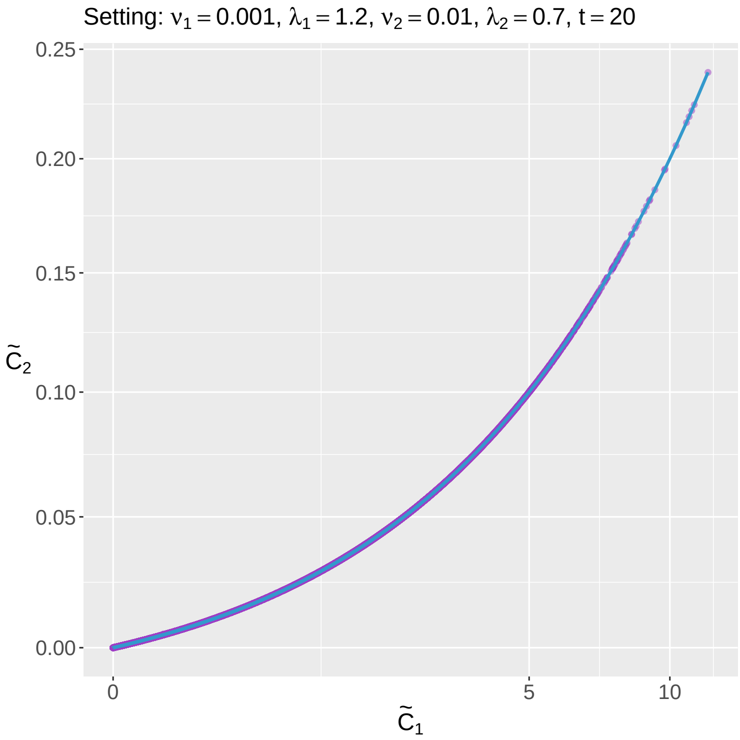

Figure S10: Comparison between the simulated cell numbers and the approximated log-normal distribution for the case $\lambda_1 = 1.2 > 0.7 = \lambda_2$. The cell numbers have been converted to their time-independent counterparts based on Theorem 1. The purple dots are the simulated values. The blue line is the theoretical mean given in Eq. (67). The blue region corresponds to the interval between the 95% and 5% quantiles of the log-normal distribution defined in Eq. (84).

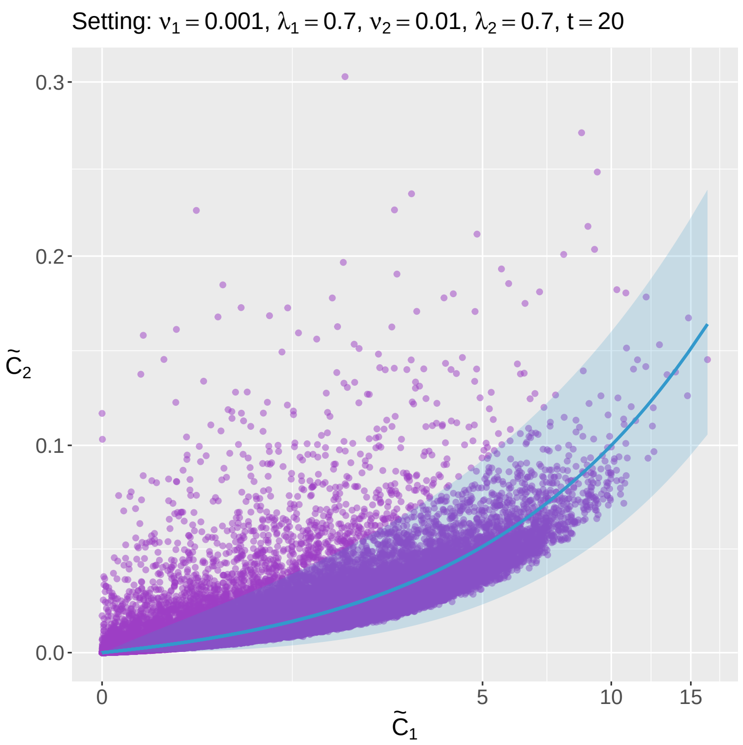

Figure S11: Comparison between the simulated cell numbers and the approximated log-normal distribution for the case $\lambda_1 = 0.7 = \lambda_2$. The cell numbers have been converted to their time-independent counterparts based on Theorem 1. The purple dots are the simulated values. The blue line is the theoretical mean given in Eq. (67). The blue region corresponds to the interval between the 95% and 5% quantiles of the log-normal distribution defined in Eq. (84).

##### D.2 Tree generating process

**Algorithm 1:** Tree generation using the tau-leaping method

**Input:**

$n$: Number of mutations
$N$: Number of trees to sample
$\boldsymbol{\mu} = (\mu_i)_{i \in [n]}$: Mutation rates of individual mutational events
$F$: An $n$-by-$n$ fitness parameter matrix
$\beta$: A common death rate (*e.g.* 1.0)
$C_0$: Static wild-type population size (*e.g.* $10^5$ cells)
$C_{\text{seq}}$: Number of cells being sequenced (*e.g.* $10^4$ cells)
$C_{\text{sampling}}$: Scale of the number of cells at sampling (*e.g.* $10^9$ cells)
$R$: Tree expansion rule (*e.g.* ISA, no ISA, etc. )
$\tau$: Tau-leaping step size (*e.g.* 0.001)
$t_{\max}$: Maximum runtime allowed (*e.g.* 100)

**Output:**

$\{\mathscr{T}_1, \ldots, \mathscr{T}_N\}$: A set of $N$ mutation trees

**for** $i = 1, \ldots, N$ **do**
  `/* Initialization */`
  Initialize time: $t = 0$
  Initialize tree $\mathscr{T}_i$ with the root $v_0$: set $C_{v_0}(t) = C_0$
  Initialize the sampling event $S(t) = 0$

  `/* Gillespie loop */`
  **while** $S(t) = 0$ *and* $t < t_{\max}$ **do**
    `/* Tree expansion */`
    **for** $v \in V(\mathscr{T}_i)$ **do**
      **if** $C_v(t) > 0$ **then**
        Expand $\mathscr{T}_i$ based on the tree expansion rule $R$: $\mathscr{T}_i = \mathscr{T}_i \cup \text{ch}(v)$

    `/* Tau-leaping */`
    Calculate propensities associated with all reactions for $v \in \mathscr{T}_i$ and $S$
    Calculate the number of reactions to occur in time step $\tau$
    Update $S$ and cell counts for $v \in \mathscr{T}_i$
    Update time $t = t + \tau$

  `/* Sequencing of cells */`
  Sample without replacement $C_{\text{seq}}$ cells from the tumor cell population:

$$(\kappa_v)_{v \in \mathscr{T}_i, v \neq v_0} \sim \text{Multivariate-Hypergeometric}(\{C_v(t)\}_{v \in \mathscr{T}_i, v \neq v_0}, C_{\text{seq}}) \tag{88}$$

  `/* Tree truncation */`
  Recursively remove the leaf nodes with zero sequenced cells from $\mathscr{T}_i$

Output $\{\mathscr{T}_1, \ldots, \mathscr{T}_N\}$

#### D.3 Ground-truth parameters for tumor mutation tree generation

We use the following parameters to generate tumor mutation trees:

- Tree expansion rule $R$: we allow for parallel mutations to be observed in different lineages, consistent with the AML dataset;
- Number of wild-type cells $C_0$: we set $C_0 = 10^5$, which is the average HSC reserve size;
- Scaling factor of the tumor size at sampling $C_{\text{sampling}}$: we choose $C_{\text{sampling}} = 10^8$ to limit computational resources required for simulation;
- Number of sequenced cells $C_{\text{seq}}$: we set $C_{\text{seq}} = 10^4$ to be consistent with the AML dataset;
- Maximum observation time $t_{\max}$: we set $t_{\max} = 100$ to be consistent with the AML dataset;
- Mutation rates $\boldsymbol{\mu} = (\mu_i)_{i \in [n]}$: all mutation rates are set to $3 \times 10^{-7}$ to limit computational resources required for simulation;
- Common death rate $\beta$: we set $\beta = 1$ to be consistent with the AML dataset;
- Fitness matrix $F$: we sample each diagonal entry from a spike-and-slab prior [6], with a 50% probability of being zero and a 50% probability of being drawn from a log-normal distribution with targeted mean and standard deviation being 0.12 and 0.03 respectively. Then, we randomly sample 50% of them and set them to positive. The off-diagonal entries are sampled from another spike-and-slab prior, with a 50% probability of being zero and a 50% probability of being drawn from a $\mathcal{N}(0, 0.25)$ distribution. These values are chosen such that in combination with $C_{\text{sampling}} = 10^8$ and $\mu_i = 3 \times 10^{-7}$, the distributions of tree sizes and the lifetime risk are consistent with the AML dataset.

#### D.4 Details on method execution

For FiTree, we assign a prior $\mathcal{N}(0, 0.01)$ to all entries in the fitness matrix $F$. Additionally, we impose a normal prior centered around the true lifetime risk with standard deviation 0.001 on the model lifetime risk. Additionally, we impose a normal prior on the model lifetime risk, centered around the true lifetime risk with a standard deviation of 0.001. To align with the analysis of the AML dataset, we first count the occurrences of each genotype across the cohort of trees. Mutations in genotypes that appear in fewer than five trees are excluded from inference by applying masks to their off-diagonal entries, thereby reducing the complexity associated with high dimensionality. The Markov Chain Monte Carlo (MCMC) sampling procedure is conducted with 1000 tuning samples and 1000 draws across 12 independent chains. This setting is used to create the plots in the main text. In Figure S12 we demonstrate that excluding infrequently mutated genes from inference does not introduce significant bias in the estimated fitness landscape. Furthermore, without the informative prior on the lifetime risk, the fitness effects are prone to inaccuracies Figure S13.

SCIFIL [7] is specifically designed for single-patient analysis; therefore, we apply it to each individual tree to estimate the fitness effect of each subclone. The fitness effects obtained from all trees are then averaged to derive the final estimates.

Similarly, fitclone [8] is also designed for single-patient analysis and relies on longitudinal samples. For each tree, we create two time points: the first at $t = 0$, representing the presence of only the wild-type population, and the second at the time of diagnosis. We then estimate the fitness for each tree individually and average the results across all trees to obtain the final estimates. Since fitclone is a Bayesian approach, we run it with 1000 tuning samples and 1000 draws.

The diffusion approximation method in [9] requires variant allele frequencies (VAFs) as input, which we calculate as

$$\text{VAF} = \frac{\sum \text{sizes of subclones with the variant}}{2 \times \text{tumor size}}. \tag{89}$$

Then, we perform maximum likelihood estimation of the fitness using the normalized version of density given in Eq.1 of the paper.

#### D.5 Additional figures

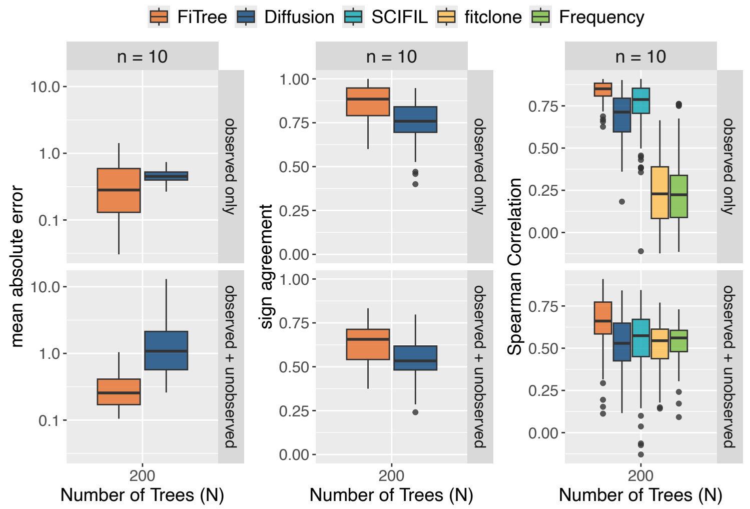

Figure S12: Performance comparison of FiTree with state-of-the-art methods in learning fitness landscapes on simulated data. Boxplots display the mean absolute errors (a), sign agreement (b), and Spearman correlations (c) across 100 simulation runs for $n = 10$, $N = 200$, and evaluation scope (observed subclones only or including unobserved subclones). For FiTree and fitclone, each point is derived from the posterior medians, while for other methods, points correspond to maximum likelihood estimates. Boxes represent the interquartile range (IQR) with medians marked inside, whiskers extend to 1.5 times the IQR, and outliers are shown as individual points. This plot differs from the main text that we did not apply masks on less frequently observed mutations in the FiTree inference.

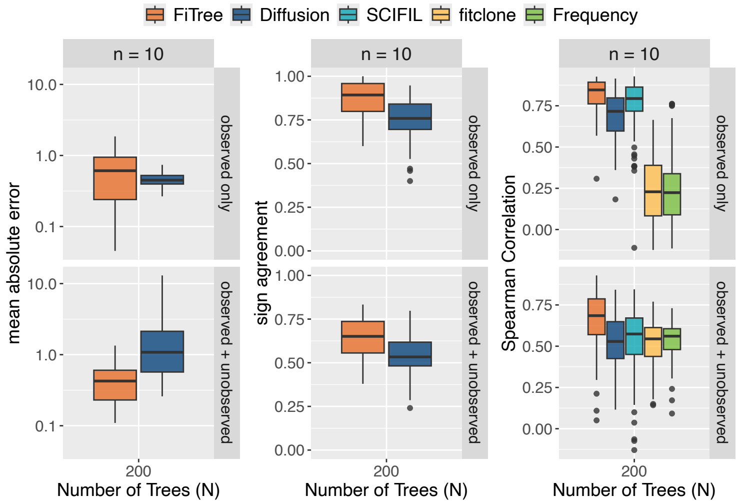

Figure S13: Performance comparison of FiTree with state-of-the-art methods in learning fitness landscapes on simulated data. Boxplots display the mean absolute errors (a), sign agreement (b), and Spearman correlations (c) across 100 simulation runs for $n = 10$, $N = 200$, and evaluation scope (observed subclones only or including unobserved subclones). For FiTree and fitclone, each point is derived from the posterior medians, while for other methods, points correspond to maximum likelihood estimates. Boxes represent the interquartile range (IQR) with medians marked inside, whiskers extend to 1.5 times the IQR, and outliers are shown as individual points. This plot differs from the main text that we neither apply masks on less frequently observed mutations nor use informative prior on the model lifetime risk in the FiTree inference.

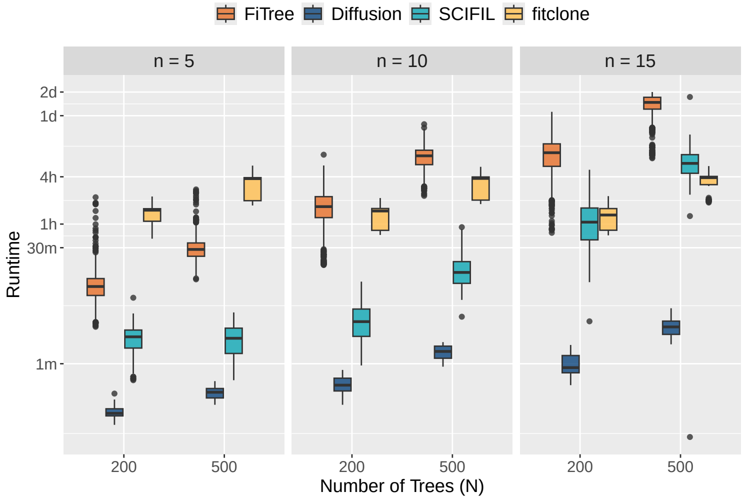

Figure S14: Runtime comparison for all methods in simulation studies.

### E Application to AML mutation trees

#### E.1 Data preparation and static parameter estimation

Following [10], we summarize somatic variants at the gene level and observe that many mutations appear in parallel branches of the same tree, *e.g. FLT3*, *PTPN11*, *NRAS*, etc. The tree expansion rule $R$ therefore allows for parallel mutations. While the observed point mutations within the same gene may have different fitness effects, this aggregation enhances the statistical power to detect non-random patterns, as gene-level events occur more frequently than individual variant-level events (Figure S15).

The total number of sequenced cells $C_{\text{seq}}$ is available for each patient. To estimate the total number of leukemia cells $C_{\text{tumor}}$, we leverage the clinical data of the cohort [11]:

$$\begin{aligned} &\text{PB\_WBC} \times \text{PB\_Blast} \times \text{Average adult blood volume} \\ &+ \text{BM\_Blast} \times \text{Average adult bone marrow mononuclear cell count} \end{aligned} \tag{90}$$

- PB_WBC: white blood cell counts in peripheral blood (in $1 \times 10^3$/µL)
- PB_Blast: blasts in peripheral blood (%)
- BM_Blast: blasts in bone marrow (%)
- Average adult blood volume: $5 \times 10^6$ µL [12]
- Average adult bone marrow mononuclear cell count: $7.53 \times 10^{11}$ [13]

We then estimate the subclone sizes by multiplying the total number of leukemia cells by the sample proportions, assuming that they accurately reflect the true cell compositions in patients. The population of human hematopoietic stem cells (HSCs) grows from birth through adolescence and then stabilizes, resulting in a relatively static population with an average of approximately $C_0 = 10^5$ wild-type cells [14, 15], which we model as the root in our framework. We assume that evolution begins once the HSC reserve fills, average around 18 years [14], which is subtracted from the patients' ages at diagnosis to get the sampling times $\boldsymbol{t}_s$ of the tumors. The maximum observation time $t_{\max}$ is then set as the maximum of the sampling times. We take the common death rate $\beta = 1.0$ to match roughly the average birth rate of HSCs [9, 14]. The lifetime risk of AML is approximately 0.5% according to the [16].

To estimate the mutation rates $\boldsymbol{\mu}$ for the 31 genes, we first calculate the rates for the corresponding 530 somatic variants using the method in [9]. For each single-nucleotide variant, we query its trinucleotide context, defined as the pyrimidine base change along with the neighboring 5' and 3' bases [15], *e.g. SRSF2* p.P95L (C[C>T]C). Next, we look up its haploid trinucleotide-context-site-specific mutation rate per year from Table S4 in [9]. For all other types of variants, we assign an average mutation rate of $2.7 \times 10^{-9}$ per bp per year, based on [15]. Finally, we sum the rates of all variants belonging to the same gene to obtain the gene-level mutation rates (Table S3). Note that we do not explicitly distinguish between tumor suppressor genes (TSGs) and oncogenes, nor do we include all mutational hotspots within those genes. On one hand, variants in TSGs do not need to be homozygous to inactivate the gene, *e.g.* mutations in codon R882 in *DNMT3A* [17]. In fact, many of them were heterozygous and already functional [11]. Additionally, loss of heterozygosity (LOH) often occurs relatively quickly after the gain of a point mutation in TSGs [18]. Therefore, the benefit of modeling the rates of TSGs differently by accounting for LOH is not significant in our case. On the other hand, the fitness values we estimate for this dataset are averages across the observed variants within genes, which is already a strong assumption as the variants may vary in fitness themselves, let alone generalizing the effects to other variants of the same genes. The median number of observed variants is 5.5 per TSG and 3.5 per oncogene, which aligns with the conventional belief that there are more positions to inactivate a TSG than to activate an oncogene [19].

We perform MCMC sampling with 1500 tuning steps followed by 1000 draws across 24 independent chains. A normal prior $\mathcal{N}(0, 0.01)$ is assigned to all elements of the fitness matrix $F$. We apply a normal prior centered at the true lifetime risk of 0.5% with a standard deviation of $10^{-8}$. Mutations observed in fewer than three trees are excluded from the inference process.

| Gene | Estimated Mutation Rate |
|---|---|
| ASXL1 | $2.430 \times 10^{-8}$ |
| BCOR | $1.350 \times 10^{-8}$ |
| CBL | $1.630 \times 10^{-8}$ |
| CSF3R | $2.700 \times 10^{-9}$ |
| DNMT3A | $1.138 \times 10^{-7}$ |
| ETV6 | $2.700 \times 10^{-9}$ |
| EZH2 | $5.054 \times 10^{-8}$ |
| FLT3 | $1.297 \times 10^{-7}$ |
| GATA2 | $6.238 \times 10^{-9}$ |
| IDH1 | $4.527 \times 10^{-8}$ |
| IDH2 | $1.853 \times 10^{-8}$ |
| JAK2 | $1.332 \times 10^{-9}$ |
| KIT | $4.533 \times 10^{-10}$ |
| KRAS | $4.871 \times 10^{-8}$ |
| MPL | $1.415 \times 10^{-8}$ |
| MYC | $1.577 \times 10^{-8}$ |
| NPM1 | $1.350 \times 10^{-8}$ |
| NRAS | $2.006 \times 10^{-8}$ |
| PHF6 | $1.470 \times 10^{-8}$ |
| PPM1D | $8.100 \times 10^{-9}$ |
| PTPN11 | $1.674 \times 10^{-8}$ |
| RUNX1 | $1.160 \times 10^{-7}$ |
| SETBP1 | $1.200 \times 10^{-8}$ |
| SF3B1 | $2.242 \times 10^{-9}$ |
| SMC3 | $4.833 \times 10^{-10}$ |
| SRSF2 | $4.372 \times 10^{-9}$ |
| STAG2 | $8.100 \times 10^{-9}$ |
| TET2 | $6.022 \times 10^{-8}$ |
| TP53 | $5.110 \times 10^{-8}$ |
| U2AF1 | $2.698 \times 10^{-9}$ |
| WT1 | $9.685 \times 10^{-8}$ |

Table S3: Gene-specific mutation rates

#### E.2 Comparison with alternative methods

In addition to FiTree, we applied SCIFIL, fitclone, and the diffusion approximation method in [9] to the AML dataset. We use the same procedure as described in Section D.4. The estimated fitness values for the top 20 most abundant genotypes averaged across all patients are shown in Supplementary Table S4. For comparability, all fitness estimates are converted to represent the relative increase from the wild-type division rate (*e.g.* a value of 0.3 indicates a 30% increase over the wild-type rate). Estimates from fitclone are reported in their original form, as fitclone does not adopt the standard fitness definition and its estimates are not directly comparable. The runtimes are displayed in Supplementary Table S5.

| Genotype | FiTree | Diffusion | SCIFIL | fitclone |
|---|---|---|---|---|
| NRAS | 0.357689 | 2.114386 | $5.63 \times 10^{-4}$ | −0.019869 |
| TP53 | 0.325165 | 9.654418 | $4.40 \times 10^{-4}$ | −0.046272 |
| IDH2 | 0.289536 | 0.727121 | $4.62 \times 10^{-4}$ | −0.044756 |
| DNMT3A | 0.247691 | 0.374935 | $3.87 \times 10^{-4}$ | −0.036987 |
| NPM1 | 0.278669 | 0.392306 | $4.47 \times 10^{-4}$ | −0.068447 |
| FLT3 | 0.277173 | 1.670385 | $4.50 \times 10^{-4}$ | −0.044658 |
| TET2 | 0.246564 | 0.361457 | $3.52 \times 10^{-4}$ | −0.019579 |
| SRSF2 | 0.248105 | 0.162225 | $3.99 \times 10^{-4}$ | −0.063024 |
| FLT3, DNMT3A | 0.229587 | 0.424869 | $8.10 \times 10^{-4}$ | −0.056460 |
| FLT3, NPM1 | 0.256709 | 0.350805 | $9.19 \times 10^{-4}$ | 0.000289 |
| DNMT3A, IDH2, SRSF2 | 1.173466 | 0.101761 | $1.41 \times 10^{-3}$ | −0.009316 |
| SF3B1 | 0.247487 | 0.160412 | $4.44 \times 10^{-4}$ | −0.044560 |
| WT1 | 0.263490 | 9.033183 | $4.38 \times 10^{-4}$ | −0.040527 |
| KRAS | 0.228479 | 0.130606 | $3.67 \times 10^{-4}$ | −0.055506 |
| NPM1, PTPN11 | 0.951057 | 0.326332 | $9.25 \times 10^{-4}$ | −0.018124 |
| DNMT3A, IDH2 | 0.238634 | 0.090128 | $7.95 \times 10^{-4}$ | −0.047589 |
| DNMT3A, TP53 | 0.222163 | 0.118064 | $9.53 \times 10^{-4}$ | 0.006947 |
| RUNX1 | 0.237887 | 0.389262 | $3.76 \times 10^{-4}$ | −0.016126 |
| DNMT3A, NRAS | 0.150791 | 0.164689 | $7.89 \times 10^{-4}$ | 0.004960 |
| GATA2, MYC | 0.302893 | 3.909406 | $1.24 \times 10^{-3}$ | −0.085379 |

Table S4: Fitness estimates for the top 20 most abundant genotypes in the AML cohort.

| Method | Runtime |
|---|---|
| FiTree | 9h 37m |
| Diffusion | 47s |
| SCIFIL | 38m |
| fitclone | 47m |

Table S5: Runtimes for different fitness inference methods applied to the AML dataset. Note that we used parallel computing (123 nodes for 123 trees) when running fitclone, whereas other methods were executed sequentially through the trees without parallelization. Parallel computing is not a built-in feature of fitclone.

##### E.3 Additional figures

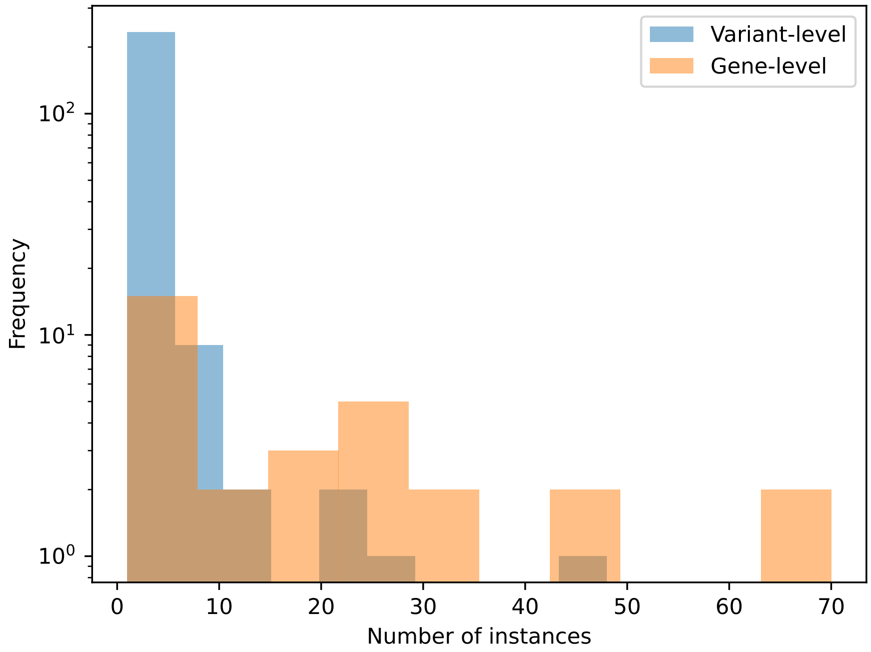

Figure S15: Distributions of mutational events at the variant level and the gene level in the AML dataset.

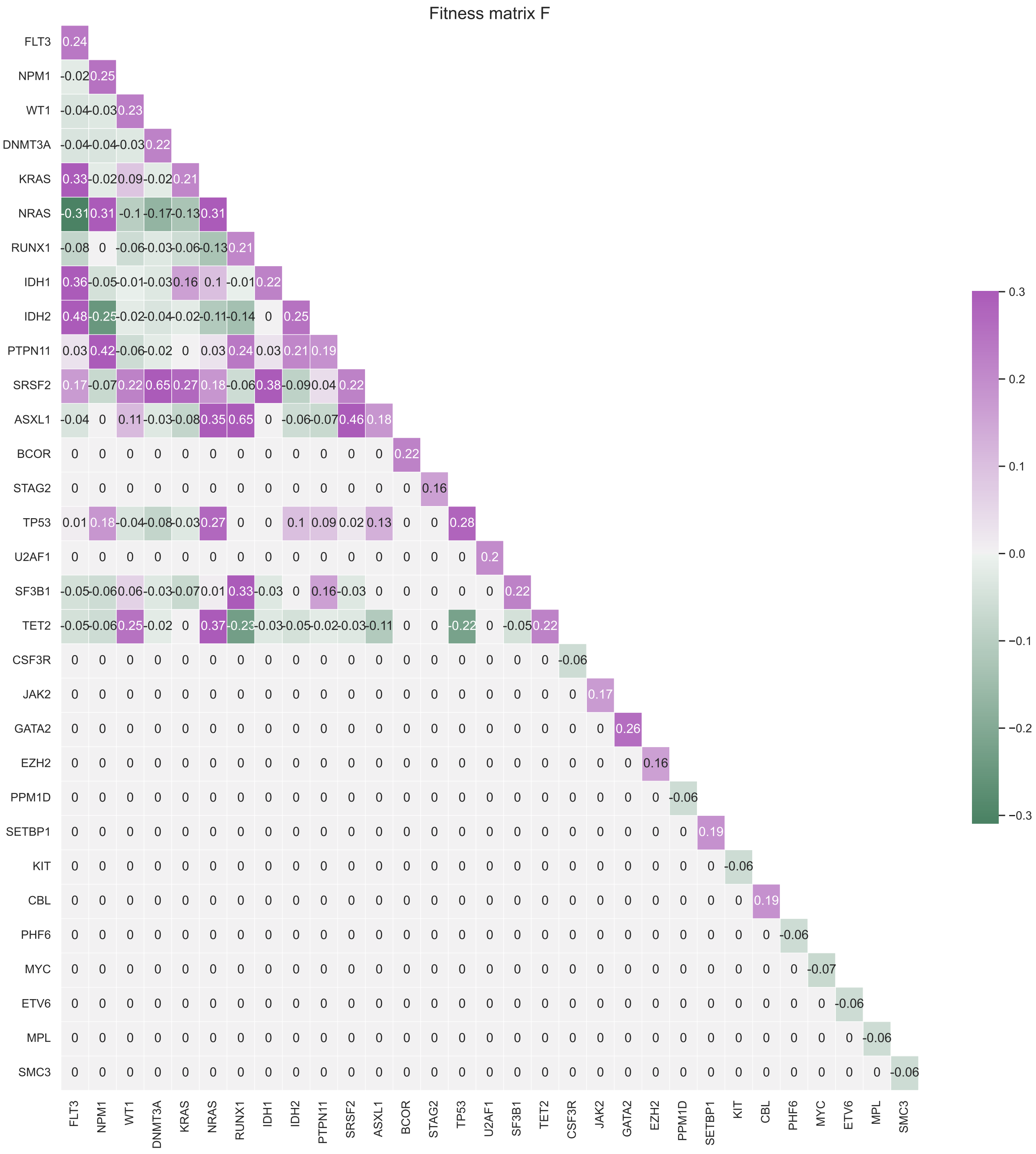

Figure S16: Posterior median of the fitness matrix for the AML dataset.

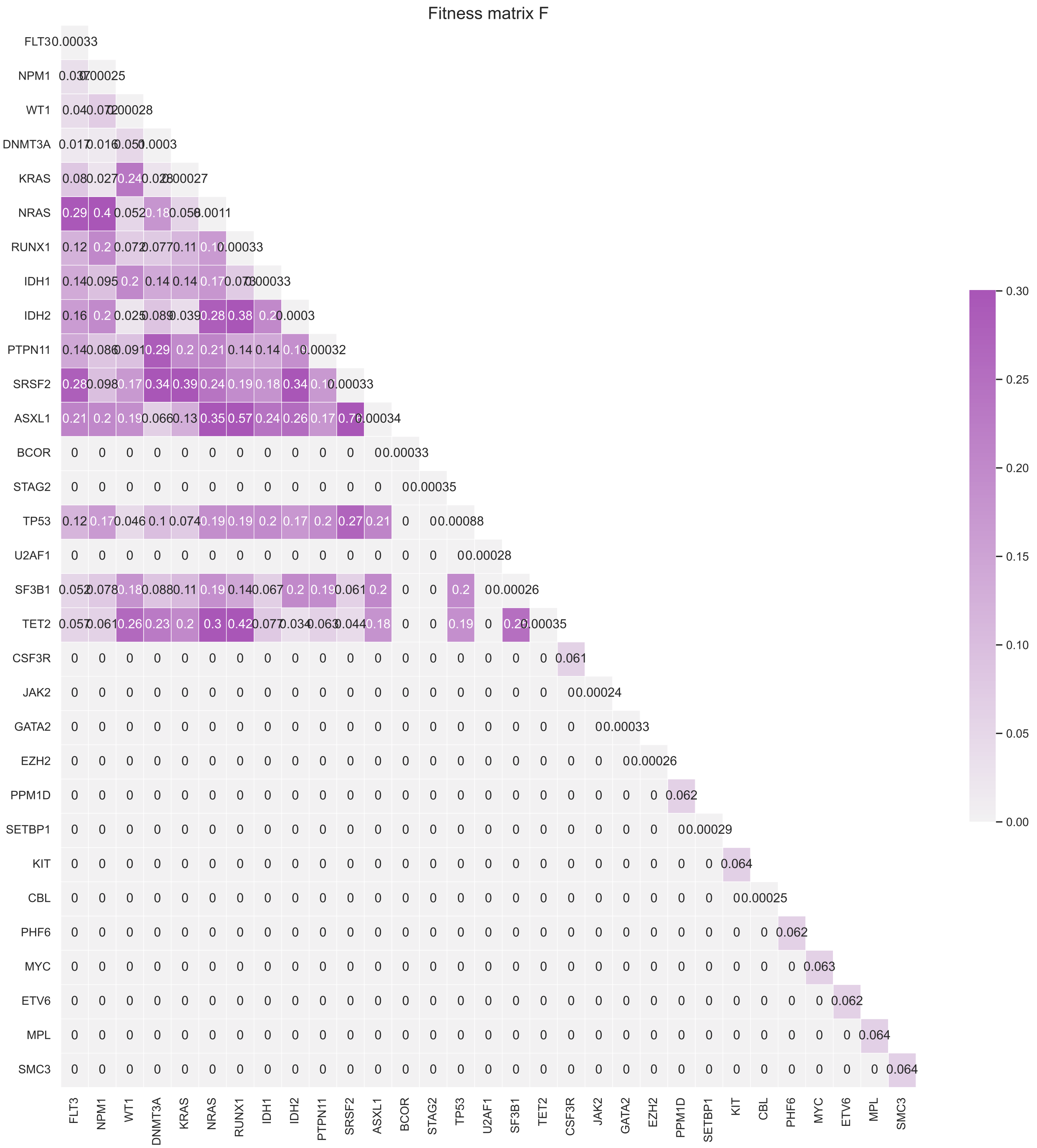

Figure S17: Standard deviation of the fitness matrix for the AML dataset.
